# Linking Systemic Endotoxin Exposure to Retinal Microglia Migration Through Mathematical Modeling

**DOI:** 10.64898/2026.06.30.735687

**Authors:** Tristen M. Jackson, Tyler Cassidy, Samantha J. Dando, Adrianne L. Jenner

## Abstract

Microglia are the resident immune cells of the central nervous system (CNS), including the brain, spinal cord, and retina, where they serve as the first line of defense against infection and inflammation. Dysregulated microglia activity has been implicated in vision-threatening diseases, highlighting the need to understand how retinal microglia respond to inflammatory stimuli. Importantly, acute inflammation induces substantial redistribution of microglia across retinal layers, yet the mechanisms governing this migration remain poorly understood. Here, we develop the first mathematical model of retinal microglia migration during inflammation to determine how inflammatory exposure, administration route, and species-specific pharmacokinetics shape redistribution dynamics across the retina. The model couples lipopolysaccharide (LPS) pharmacokinetics with microglia migration between the outer plexiform layer (OPL), inner plexiform layer (IPL), and ganglion cell layer/nerve fiber layer (GCL/NFL). Model parameters are calibrated to retinal microglia density measurements from mice following LPS (bacterial endotoxin) challenge, before extending the framework to rats and rhesus macaques to investigate species-specific responses. Simulations also compare how administration route, i.e. intravenous or intraperitoneal injections, alter retinal LPS exposure and subsequent microglia redistribution. Our results suggest that redistribution patterns are driven primarily by LPS delivery route and species-specific pharmacokinetics, rather than the initial microglia distribution across retinal layers. Together, these findings provide new insight into immune cell reorganization in the inflamed retina and demonstrate how mechanistic mathematical modeling can be adapted across experimental designs, administration routes, and animal species.

## 1 Introduction

Microglia are the resident immune cells of the central nervous system (CNS) and serve as its first line of defense against infection and inflammation. Their exclusivity reflects the immune privilege of the CNS [1], a property characterized by tightly regulated tissue barriers and mechanisms of the brain, spinal cord, and retina that safeguard the neural environment [2]. Naturally, microglia perform many of the same functions as other macrophages, including phagocytosis and cytokine secretion, but also perform specialized functions in the retina, including contributing to visual processing through modulation of synaptic activity and photoreceptor surveillance [3–5]. Unfortunately, increasing evidence implicates microglia dysregulation in the pathogenesis of vision-threatening diseases [6–8].

Systemic inflammation induces sequential responses in the peripheral immune system and CNS. One early development is increased permeability of the blood–brain and blood–retina barriers, enabling peripheral inflammatory signals to reach the CNS and initiate neuroinflammation [9, 10]. As a result, retinal microglia redistribute across the retinal layers as they mobilize to respond to inflammatory cues [11, 12]. Under homeostatic conditions, retinal microglia occupy three main layers: the outer plexiform layer (OPL), inner plexiform layer (IPL), and ganglion cell layer (GCL), see Figure 1. Following acute systemic inflammation, microglia are thought to migrate away from the OPL and IPL toward the vitreal aspect of the retina, particularly the highly vascularized GCL and NFL, where inflammatory cytokines such as TNF-α, IL-6, and IL-8 are more concentrated [3, 13–16]. Migration toward the subretinal space is also observed in other models of neural insult, such as laser-induced injury and photoreceptor degeneration.

**Figure 1:**
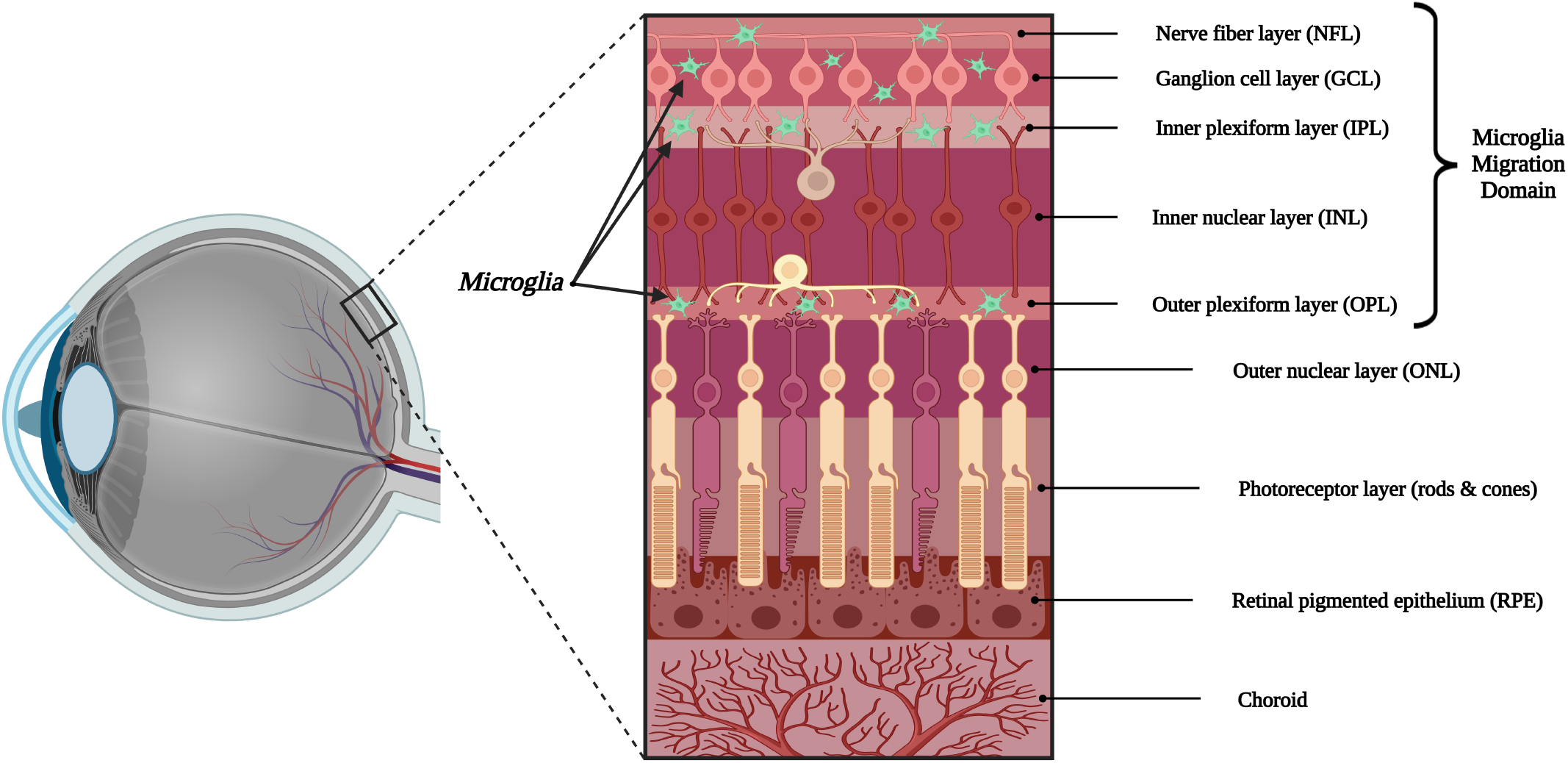
Diagrammatic representation of retinal layers and modeled microglia migration domain. The retinal architecture with major anatomical layers, highlighting the outer plexiform layer (OPL), inner plexiform layer (IPL), and ganglion cell layer/nerve fiber layer (GCL/NFL), where retinal microglia (green) reside and redistribute in response to systemic inflammation. In this study, we model the migration of microglia specifically within these layers.

Layer-specific microglia densities have been quantified in a variety of contexts using immunostaining & fluorescent imaging, and *in vivo* imaging, including under control or untreated conditions [3, 17, 18], ocular hypertension [19], optic nerve injury [11, 20], and microglia depletion [21]. Notably, Dando *et al*. [15] quantified layer-specific microglia densities at 2, 24, and 48 hours after acute systemic LPS challenge in mice, providing the most quantitative temporal characterization to date of microglia redistribution across retinal layers during inflammation. Although these studies establish a phenomenological understanding of microglia migration within the retina during inflammation, the mechanistic connection between inflammatory intensity and layer-specific microglia redistribution has not yet been quantified. Quantitatively understanding this is important for the study of retinal pathologies, with one example being that microglia migration has been shown to drive photoreceptor degeneration in a mouse model of retinitis pigmentosa [4,22].

Mechanistic mathematical modeling offers a clear opportunity to connect inflammatory intensity with layer-specific microglia redistribution by extrapolating from existing data through a series of minimal biological inferences. Its use has proliferated in peripheral immunology [23–29], yet applications to neuroinflammation and ocular inflammation remain comparatively rare [30]. For example, agent-based models have been developed to study microglia in neuro-degenerative diseases, such as Alzheimer’s disease [31–33]. Whereas, ordinary differential equations (ODEs) have been used to study microglia metabolism [34]. Other related studies have modeled microglia *M*_1_ and *M*_2_ phenotype dynamics in the ischemic penumbra following stroke [35, 36]. While these studies capture temporal changes in microglia density, their applications are to specific diseases, and not generalizable to any acute inflammatory stimuli.

In the context of the retina, a small but informative collection of ODE models has been developed to study oxygen distribution across retinal layers [37–39]. Though these models do not address immunological processes, they provide a valuable proof of concept for quantitatively describing translayer movement and compartmental structure within the retina. Related work has also used mathematical modeling to study the emergence of microglia spacing in the developing retina [40]. Although focused on developmental organization rather than inflammatory responses, this study succeeded at providing a spatiotemporal description of microglia location.

In the present work, we develop the first mathematical model of retinal inflammation with a focus on the migration of microglia across the upper most layers of the retina (see Figure 1). The model is implemented as a system of mechanistic ordinary differential equations (ODEs) describing microglia migration between the OPL, IPL and GCL/NFL during acute inflammation, calibrated initially to the experimental data of Dando *et al*. [15]. The model is composed of two interacting subsystems. The first captures the distribution and clearance of exogenously administered LPS, and the second describes LPS-mediated microglia migration between the OPL, IPL, and GCL/NFL. To investigate the impact of LPS delivery route and species, the model is reparameterized to a range of different data sets enabling predictions of how microglia migration dynamics change under different LPS doses, routes of administration, and experimental animal species.

## 2 Methods

### 2.1 Details of Experimental Measurements

We utilized data from a series of experimental set-ups and model organisms to fit parameters in our model. Unless otherwise specified, data were extracted from published figures using WebPlotDigitizer [41].

#### 2.1.1 Intraperitoneal LPS Pharmacokinetics in Mice

Huang *et al*. administered a single intraperitoneal (IP) injection of lipopolysaccharide (LPS) derived from *E. coli* O111:B4 to female C57BL/6 mice [42]. Mice received doses of 0, 0.5, or 1.0 mg/kg LPS, and plasma samples were collected at 0.5, 2, 4, 12, and 24 hours post-injection. Circulating LPS concentrations were quantified from plasma using a Limulus amebocyte lysate (LAL) chromogenic endpoint assay.

#### 2.1.2 Time-course Retinal Microglia Densities in Mice

Dando *et al*. [15] delivered LPS from *E. coli* OIII:B4 at 9 mg/kg to BALB/c *Cx3cr1*^*gfp/gfp*^ mice. Microglia densities were quantified separately within the OPL, IPL, and NFL/GCL at multiple time points following LPS administration (control, 2 h, 24 h, and 48 h) via confocal microscopy. Densities in the OPL, IPL, and GCL/NFL were quantified in units of cells per mm^2^.

#### 2.1.3 Intravenous LPS Pharmacokinetics in Mice

Yao *et al*. administered a single intravenous (IV) infusion of rough LPS via the tail vein to adult male BALB/c and C57BL/6 mice [43]. LPS was derived from *Escherichia coli* K-12 and *Salmonella enterica* serovar Typhimurium and delivered at doses of approximately 7–10 *µ*g per mouse. Blood samples were collected at 0.5, 2.5, 5, 15, and 30 minutes post-administration. Circulating LPS concentrations were normalized by body weight prior to reporting.

#### 2.1.4 Intraperitoneal LPS Pharmacokinetics in Rats

Hutter & Kim administered a single IP injection of 200 *µ*g/kg lipopolysaccharide (LPS) derived from *Salmonella typhimurium* to adult Sprague–Dawley rats [44]. Blood samples were collected at 5, 15, and 30 minutes, and at 1, 2, 4, 6, 8, 12, 18, 24, and 48 hours post-administration. LPS was dual-labeled with glucosamine and fatty acid residues, enabling independent quantification of LPS components. Because fatty acid residues are incorporated into long-lived lipid pools following LPS cleavage, plasma glucosamine concentrations were used as the primary measure of circulating LPS over time.

#### 2.1.5 Intravenous LPS Pharmacokinetics in Rhesus Monkeys

Mathison *et al*. administered a single IV injection of lipopolysaccharide (LPS) derived from *Salmonella minnesota* R595 to adult male rhesus macaques (6.5–7 kg) [45]. Animals received a dose of 5 mg/kg LPS via a femoral vein catheter. Serial blood samples were collected at early time points (0.75, 2, 3, 4, and 5 minutes post-injection) to capture the rapid distribution phase, with additional samples obtained at later times (approximately 20, 180, 225, and 300 minutes post-injection) to characterize the slower systemic clearance phase.

#### 2.1.6 Homeostatic Retinal Microglia Densities in Rhesus Monkeys

Data from Singaravelu *et al*. were used to estimate baseline microglia densities in untreated, healthy, young adult rhesus macaques [18]. In this study, microglia densities were reported separately for each retinal eccentricity region (fovea, macula, and peripheral retina). Following euthanasia, retinal tissue was fluorescently stained using Iba-1 antibodies and imaged using confocal microscopy to quantify microglia densities. Because Iba-1 is not specific to microglia and also labels infiltrating myeloid cells, these measurements are interpreted as approximations of resident retinal microglia density.

### 2.2 Mathematical Models

To investigate how acute systemic inflammation drives microglia redistribution across retinal layers, we developed two related mathematical models that couple LPS pharmacokinetics with layer-specific microglia migration. These models differ in their pharmacokinetic structure, representing either intraperitoneal (IP) delivery (Figure 2A) or intravenous (IV) delivery (Figure 2B), and are each coupled to a common microglia migration subsystem (Figure 2C). Each coupled model was subsequently parameterized to two distinct experimental contexts, yielding four scenario-specific model formulations used throughout this study: IP injection in mice, IV injection in mice, IP injection in rats, and IV injection in monkeys. All model simulations, parameter estimation routines, and figure-generation code used in this study are publicly available on GitHub.

**Figure 2:**
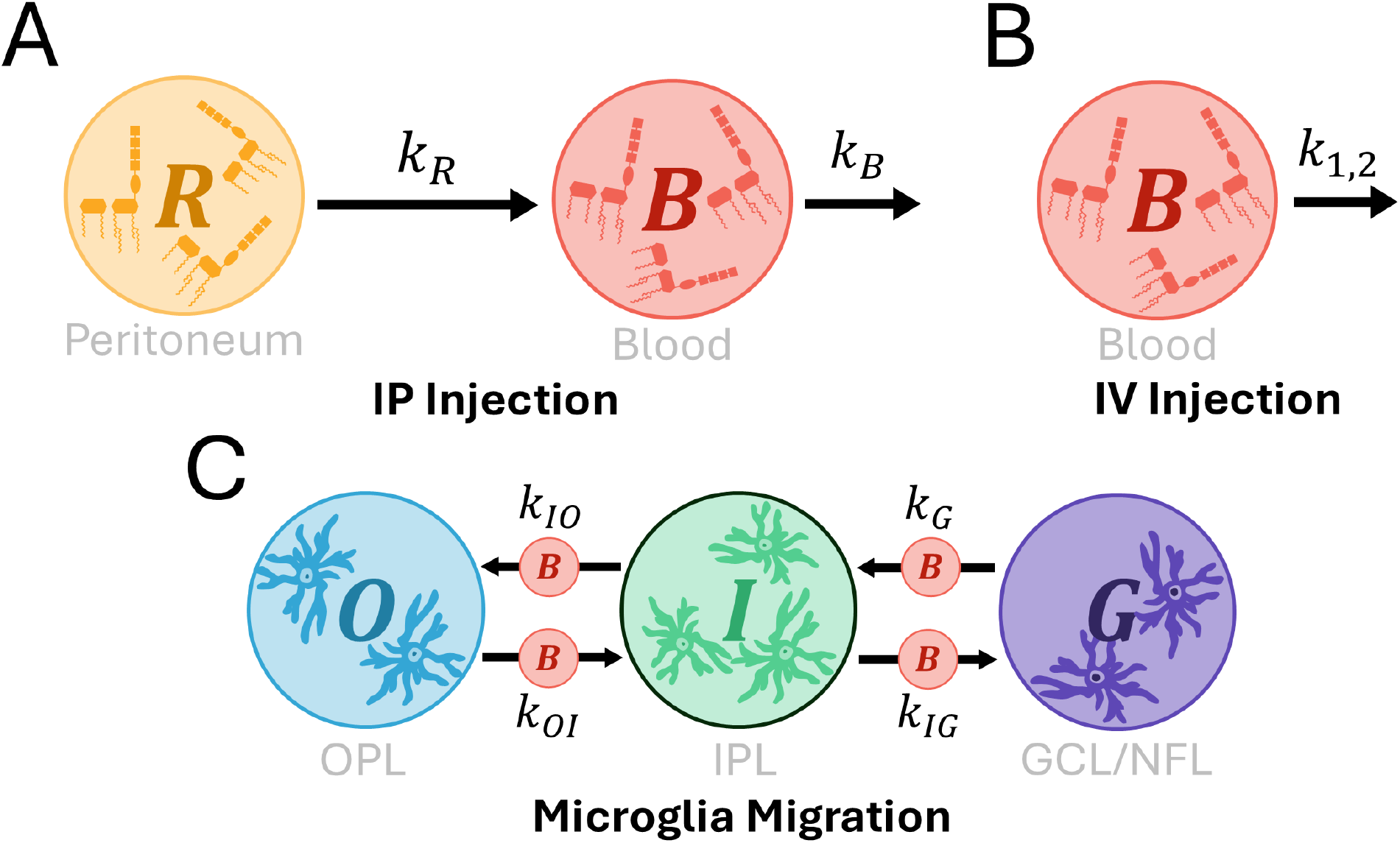
Schematic representation of the coupled mathematical models used throughout this study. **(A)** Intraperitoneal pharmacokinetic subsystem, describing the transfer of LPS from the peritoneal cavity to the bloodstream. The parameter *k*_*R*_ represents the rate of LPS transfer from the peritoneum to bloodstream, while *k*_*B*_ represents the rate of LPS clearance from the bloodstream. **(B)** Intravenous pharmacokinetic subsystem, in which LPS is introduced directly into the blood, and the peritoneal compartment is omitted. **(C)** Microglia migration subsystem, describing movement between the OPL, IPL, and GCL/NFL. Parameter *k*_*i j*_ represents migration from compartment *i* to compartment *j*, where *k*_*G*_ denotes movement from GCL/NFL to IPL. Final models were constructed by coupling either the intraperitoneal or intravenous pharmacokinetic subsystem with the microglia migration subsystem, yielding two distinct model structures.

The pharmacokinetic component describes the concentration of LPS (ng/mL) in the peritoneal cavity, *R* (*t*), and its subsequent appearance in the bloodstream, *B* (*t*) . Note that in the intravenous model, LPS is introduced directly into the blood compartment and the peritoneal variable *R* (*t*) is omitted. The microglia migration component describes the densities (cells/mm^2^) of microglia residing in the outer plexiform layer, *O*(*t*), inner plexiform layer, *I* (*t*), and ganglion cell layer/nerve fiber layer, *G*(*t*).

#### 2.2.1 Intraperitoneal LPS Subsystem

The peritoneal compartment is modeled as a first-order absorption process, representing the transfer of LPS from the peritoneal cavity into systemic circulation at rate *k*_*R*_. A bioavailable fraction *F* of peritoneal LPS enters the bloodstream, scaled by the ratio of peritoneal to blood volumes, *Vol*_*p*_ / *Vol*_*B*_. We scale by the ratio of compartment volumes to ensure that the concentration of LPS leaving the peritoneum is converted into the corresponding concentration increase in the blood compartment. Furthermore, the fraction of IP-administered LPS that reaches systemic circulation, *F*, is assumed to be independent of dose size. Once in circulation, LPS is removed from the blood through first-order clearance at rate *k*_*B*_. The resulting two-compartment subsystem is provided below, where *t* denotes time in hours:

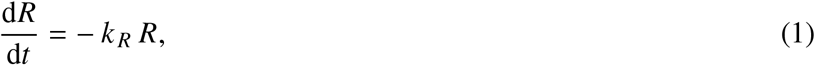

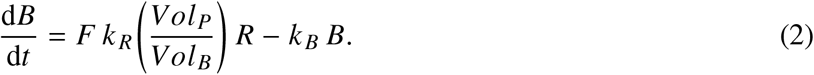

The LPS kinetics following IP injection are monophasic, which is consistent with no distribution to the tissues from the circulation. We also note, **Equations (1)-(2)** can be solved explicitly for *R*(*t*) and *B*(*t*), see Supplementary Information for their explicit form.

#### 2.2.2 Intravenous LPS Subsystem

In contrast to intraperitoneal administration, intravenous injection delivers LPS directly into systemic circulation. As a result, we chose not to explicitly model any transport in or out of the tissues. Experimental studies consistently report that circulating LPS following intravenous administration exhibits biphasic decay, reflecting a rapid initial decline and distribution to the tissues, followed by a slower terminal phase [43,46,47]. The IV data used in this paper includes densely sampled early time points which allows us to observe clear biphasic decay. Accordingly, the intravenous LPS subsystem is modeled here using a biexponential decay function, which captures both the fast and slow clearance components observed in the literature:

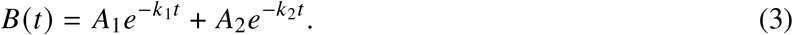

Here, *A*_1_ and *A*_2_ are fitted coefficients that weight the fast and slow exponential components, while *k*_1_ and *k*_2_ are the corresponding rate constants governing the fast and slow clearance phases respectively.

#### 2.2.3 Microglia Migration Subsystem

Microglia redistribution across retinal layers was modeled using a system of ODEs describing the densities of microglia in the OPL, *O*(*t*), IPL, *I* (*t*), and GCL/NFL, *G*(*t*). These equations represent migration between adjacent retinal layers when in the presence of circulating LPS, *B*(*t*). The final subsystem is given by:

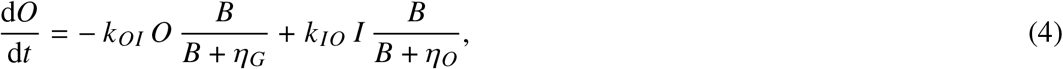

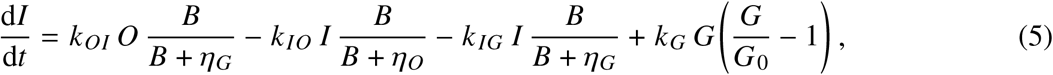

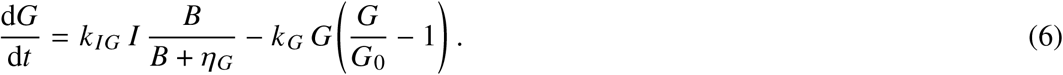

We model microglia migration between compartment *i* and *j*, using the Hill function

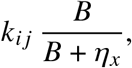

where *k*_*i j*_ denotes the baseline transition rate of microglia from compartment *i* to compartment *j*, and *η*_*x*_ is a direction-dependent half-saturation constant. The subscript *x∈* {*O, G*} reflects the direction of migration, with *x* = *O* corresponding to migration toward the OPL and *x* = *G* corresponding to migration toward the GCL/NFL. In this sense, the Hill function represents the saturating influence of circulating LPS *B*(*t*) on microglia transition rates. We assume microglia sense inflammatory signals originating from the retinal vasculature, which is primarily localized within the GCL/NFL. Consequently, the influence of circulating LPS depends on whether microglia are migrating toward or away from this vascular source. Allowing *η*_*x*_ to vary with migration direction enables the model to capture potential asymmetry in migratory responsiveness across retinal layers.

We represent migration from the GCL/NFL to the IPL using a logistic-type term of the form

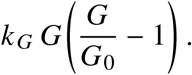

where *k*_*G*_ is the intrinsic regulation rate of microglia in the GCL/NFL. This choice reflects the assumption that, due to the proximity of the GCL/NFL to the retinal vasculature, microglia in this compartment are more tightly regulated by inflammatory signaling than those in the OPL or IPL. Consequently, once the inflammatory signal begins to weaken, microglia in the GCL/NFL are expected to return to their home-ostatic density on a shorter time scale. We capture this regulatory behavior through a logistic relaxation term that is independent of *B* (*t*), thereby enforcing return toward the baseline density *G*_0_ = *G*( 0) as the inflammatory stimulus subsides. This form reflects experimentally observed time-course data indicating more rapid normalization of microglia in the GCL/NFL compared to other retinal layers [15].

We note that **Equations 4-6** are designed to capture LPS-induced microglia redistribution over the acute experimental window of up to 48 h post-treatment. In the absence of LPS (*B* = 0), the system admits a continuum of equilibria determined by the initial conditions and is not intended to represent homeostatic dynamics. Initial densities *O* (0), *I* (0) and *G* (0) are defined from control measurements as described in the Supplementary Information. Many variations of the microglia migration subsystem (**Equations 4-6**) were evaluated and were unable to capture the data, see the Supplementary Information for details.

### 2.3 Mouse Model Parameter Estimates

The parameter estimates presented in this section reflect mouse-specific physiology and pharmacokinetics and are therefore distinct, in some cases, from the parameterizations later obtained for rats and rhesus macaques. All fitted parameters were obtained through Matlab’s lsqnonlin after log-transforming both the data and model outputs. The mathematical models obtained by coupling either the intraperitoneal pharmacokinetic subsystem (**Equations 1–2**) or the intravenous pharmacokinetic formulation (**Equation 3**) with the microglia migration subsystem (**Equations 4–6**) were structurally identifiable, as evaluated using the open-source software DAISY [48, 49] and Julia package StructuralIdentifiability.jl [50]. Despite this, limited time-series data meant that not all parameters were practically identifiable.

#### 2.3.1 Intraperitoneal LPS Subsystem

Parameters in **Equations 1-2** were either taken from literature, explicitly calculated, or fitted to data when neither of these options were possible, see **Table 1**. Because this pharmacokinetic subsystem was calibrated using experimental data from 7–8 week old female C57BL/6J mice [42], physiological volumes were selected to reflect that same age and strain. Peritoneal volume (*Vol*_*p*_) in 7-8 week old female mice was estimated at 0.1mL [51]. A small adjustment to the peritoneal volume was needed to reflect the additional 0.01mL vehicle that LPS was injected in, so the total peritoneal volume was taken as 0.1 + 0.01mL.

**Table 1.**
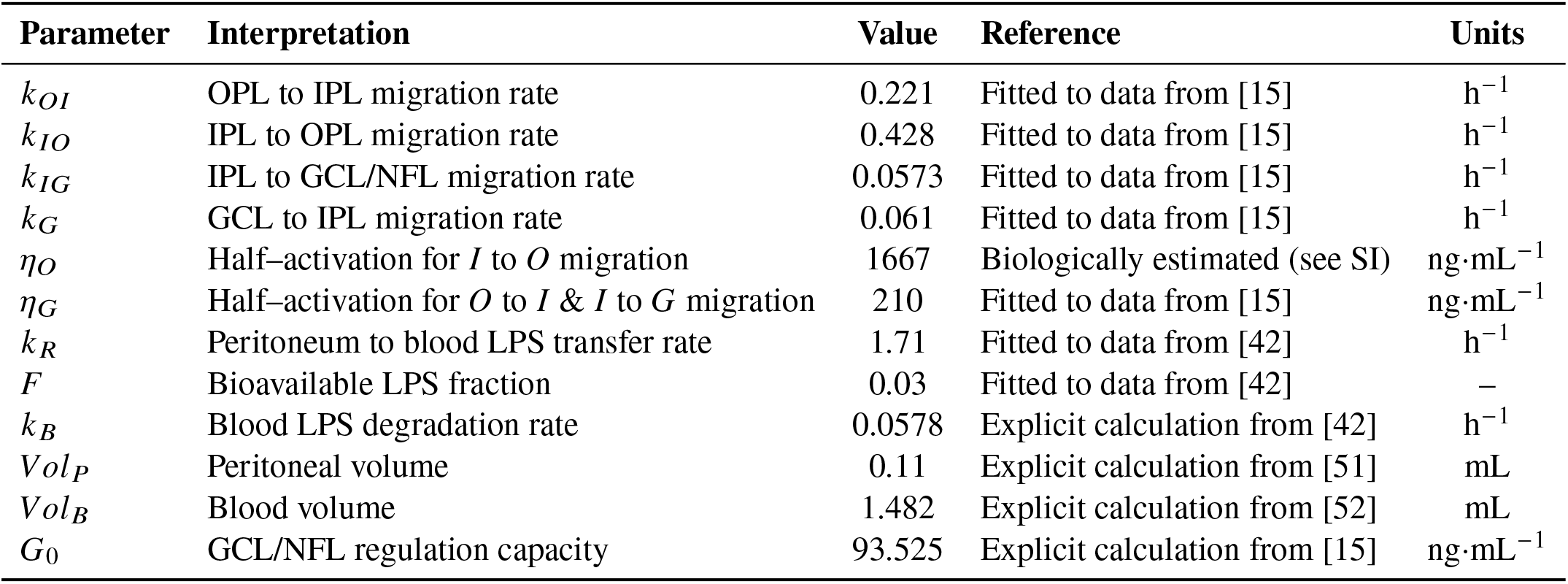
Parameter definitions, values, interpretations, and units for the intraperitoneal LPS subsystem and microglia migration model (**Equations 1-2 and 4-6)** A comparison of model simulations to *in vivo* data under this parameter regime is provided in Figure 3.

Blood volume (*Vol*_*B*_) was similarly estimated at 0.077 mL/g of body weight [52]. Since 7-8 week old female C57BL/6J mice typically weigh between 17.8 and 20.7 g [53], the midpoint of this was taken to be 19.25 grams, giving a blood volume (*Vol*_*B*_) of approximately 1.482 mL. Assuming first-order clearance of LPS from the blood following intravenous administration, the clearance rate was calculated using

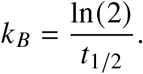

The half-life of LPS in the blood (*t*_1/2_) was estimated to be 12 hours post-injection [42], i.e. *k*_*B*_ = 0.057. The remaining parameters of *F* and *k*_*R*_ were estimated using Matlab’s lsqnonlin and log transforming the data and model. Both the 0.5 mg/kg and 1.0 mg/kg intraperitoneal LPS datasets were fitted simultaneously.

#### 2.3.2 Intravenous LPS Subsystem

Best-fit parameters were obtained by minimizing the squared error between log-transformed measured and predicted blood concentrations over the experimental time window, with amplitudes *A*_1_ and *A*_2_ constrained such that *A*_1_ + *A*_2_ equals the measured initial plasma LPS concentration. The resulting parameter estimates are provided in **Table 2**.

**Table 2.** Fitted parameter values for the intravenous LPS biexponential decay model (**Equation 3**), obtained by nonlinear least-squares fitting to plasma LPS concentration data from Yao *et al*. [43] following intravenous administration in mice.

| Parameter | Interpretation | Value | Units |
| --- | --- | --- | --- |
| $A_1$ | Fast-phase amplitude | 0.6654 | – |
| $A_2$ | Slow-phase amplitude | 0.3307 | – |
| $k_1$ | Fast clearance rate | 0.2512 | $\text{min}^{-1}$ |
| $k_2$ | Slow clearance rate | 0.0204 | $\text{min}^{-1}$ |

#### 2.3.3 Microglia Migration Subsystem

The microglia migration subsystem defined by **Equations 4 -6** is composed of six unknown parameters, including the microglia migration rates (*k*_*OI*_, *k*_*IO*_, *k*_*IG*_, *k*_*G*_) and the LPS stimulation parameters (*η*_*G*_ and *η*_*O*_). For the data in this study, *η*_*O*_ and *η*_*G*_ were not practically identifiable. As such, *η*_*O*_ was therefore estimated biologically, which resolved the identifiability issue for *η*_*G*_, and improved fits for *k*_*OI*_, *k*_*IO*_, *k*_*IG*_, and *k*_*G*_. These remaining parameters were subsequently fitted. Due to the absence of synthesized dose-response relationships for this process, we selected a representative intermediate LPS dose based on values reported in the literature. Experimental studies span a broad range of administered doses (approximately 0.1–9 mg/kg), from low-dose systemic challenges (0.1 mg/kg) to commonly used neuroinflammatory paradigms at 4–5 mg/kg and higher-dose retinal inflammatory models (9 mg/kg) [9, 15, 54, 55]. We therefore chose 4 mg/kg as both a pragmatic intermediate value and a literature standard that induces a clear but non-saturating neuroinflammatory response. We then simulated the PK subsystem at this dose and identified the corresponding peak blood LPS concentration (Figure S1). This peak value was taken as a proxy for the concentration required to induce a half-maximal migratory response in the OPL direction, i.e. *η*_*O*_. See the Supplementary Information and Figure S2 for more details.

The five remaining parameters were then fitted to the data published in Dando *et al*. [15] using nonlinear least squares fitting (*lsqnonlin* in Matlab). The model was fitted simultaneously to the individual microglia density measurements at each timepoint rather than to the mean values. Treating each replicate measurement as an independent observation allowed the fitting procedure to account for the full variability present in the dataset. These plots demonstrate clear minima in model error for each fitted parameter, indicating that the migration kinetics can be meaningfully interpreted. All parameters used in **Equations 1 - 6** are in Table 1. The initial conditions where fixed to the mean initial measurements, see Table S1.

### 2.4 Mathematical Model Evaluation Metrics

To evaluate model performance across all subsystems, goodness-of-fit was assessed using root mean square error (RMSE). Where appropriate, confidence intervals derived from parameter estimation were propagated through the model to assess trajectory uncertainty. For fitted pharmacokinetic parameters, approximate 95% confidence intervals were computed using the nonlinear least-squares covariance approximation. Trajectory bands were then obtained by re-simulating the ODE model for every combination of the upper and lower 95% confidence bounds of the fitted parameters, rather than using a delta-method approximation.

### 2.5 Mouse Model Sensitivity Analysis

The sensitivity of microglia densities in each layer at 48 hours, along with the AUC for each microglia density curve at 48 hours was assessed using partial rank correlation coefficients (PRCC). As a prerequisite for PRCC analysis, monotonic relationships between each parameter and model output were verified using scatter plots and corresponding Spearman rank correlation coefficients (Figure S6).

#### 2.5.1 Parameter Ranges

For the PRCC analysis, upper (*U*_*i*_) and lower (*L*_*i*_) bounds were specified for each parameter based on biological evidence, literature-derived physiological limits, or variability observed during model fitting. Hartveit *et al*. [51] reported that peritoneal fluid volume in adult female mice ranges from 0.05 to 0.15 mL, as 0.01 mL of LPS vehicle was administered intraperitoneally in the experiments used for model calibration, this additional volume shifts the effective range for *Vol*_*p*_ to [0.06, 0.16] mL. Mouse weights vary between 17.8 and 20.7 grams [52, 53], which corresponds to a range for *Vol*_*B*_ of [1.371, 1.594] mL, assuming blood volume for mice is 0.077 mL/g of their body weight. From the Huang *et al*. dataset [42], the 0.5 mg/kg dose of LPS gave a half-life estimate of 11.5 hours and the 1.0 mg/kg dose gave an estimate of 14.6 hours. The half-life range was then [11.5, 14.6] hours, which gives a range for *k*_*B*_ of [0.0475, 0.0603]/hour. The range of *F* and *k*_*R*_ was estimated from the 95% confidence intervals returned from the fitting, i.e. *F* was [0.0275, 0.0319] and *k*_*R*_ was [1.2677, 2.1483].

To estimate the range for *η*_*O*_, we chose a range of reasonable LPS dosages from the literature [56–58], i.e [2.0, 5.0] mg/kg, and then converted this into a range for *η*_*O*_, i.e. [834, 2,084] ng/ml (see the Supplementary Information). Since the point estimates for *k*_*IO*_, *k*_*OI*_, *k*_*IG*_, *k*_*G*_, and *η*_*G*_ were all obtained from fitting, the natural choice was to use the confidence intervals to generate a range for these parameters. Unfortunately, the 95% CIs for *k*_*OI*_ and *k*_*IO*_ are unbounded above (Figure S3). Instead, we chose “practical ranges” as the set of parameter values for which Δ*χ*^2^≤ 1, representing parameter values that are practically indistinguishable from the optimum. These ranges were then used in the PRCC sensitivity analysis. Since the Δ *χ*^2^ ≤ 1 cutoff needed to be used for *k*_*IO*_ and *k*_*OI*_, it felt natural to use this cutoff for the other parameters too (*η*_*G*_, *k*_*IG*_, and *k*_*G*_) rather than their 95% CIs.

To then explore model behavior under extreme biological scenarios and reduce potential bias from literature-derived estimates, the parameter ranges were expanded to

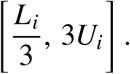

This three-fold expansion ensures that the sensitivity analysis explores a broader region of parameter space beyond experimentally observed conditions. All of the model’s parameter ranges are in **Table 3**.

**Table 3.** Parameter ranges used in the PRCC sensitivity analysis. Lower (*L*_*i*_) and upper (*U*_*i*_) bounds were first determined from biological evidence, literature-derived physiological limits, or variability observed during model fitting. These ranges were then expanded to 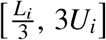 to explore a broader region of parameter space during an extended sensitivity analysis (Figure S5).

| Parameter | Sensitivity Analysis Range<br>$[L_i, U_i]$ | Extended Sensitivity Analysis Range<br>$[\frac{L_i}{3}, 3U_i]$ |
| --- | --- | --- |
| $Vol_P$ | [0.06, 0.16] | [0.02, 0.48] |
| $Vol_B$ | [1.371, 1.594] | [0.457, 4.782] |
| $k_B$ | [0.0475, 0.0603] | [0.01583, 0.1809] |
| $F$ | [0.0275, 0.0319] | [0.00917, 0.0957] |
| $k_R$ | [1.2677, 2.1483] | [0.4226, 6.4449] |
| $\eta_O$ | [834, 2084] | [278, 6252] |
| $\eta_G$ | [87.119, 372.381] | [29.040, 1117.143] |
| $k_{OI}$ | [0.1031, 0.7932] | [0.03437, 2.3796] |
| $k_{IO}$ | [0.1996, 1.3744] | [0.06653, 4.1232] |
| $k_{IG}$ | [0.0375, 0.0863] | [0.0125, 0.2589] |
| $k_G$ | [0.0365, 0.0987] | [0.01217, 0.2961] |

### 2.6 Bayesian Posterior Predictive Intervals

Standard 95% confidence intervals computed from the nonlinear least-squares fit of the migration parameters (*k*_*OI*_, *k*_*IO*_, *k*_*IG*_, *k*_*G*_) were wide, reflecting large uncertainty associated with these parameters. This uncertainty is partially addressed through our sensitivity analyses, but a clear confidence band on model trajectories was desired to showcase parameter influence on model predictions. To obtain more meaningful uncertainty quantification, we performed a Bayesian posterior predictive analysis. The residuals from the least-squares fit were used to estimate the noise variance, and all parameters were log-transformed to ensure positivity. An MCMC sampler was then run using broad, non-informative priors on both parameters and noise. Posterior samples were propagated through the full ODE model to generate posterior predictive trajectories, from which 2.5th–97.5th percentile bands were computed. These posterior predictive intervals provide a model-based quantification of trajectory uncertainty that accounts for both parameter and observational noise. The final Bayesian Predictive Posterior Intervals are provided in Figure 3.

**Figure 3:**
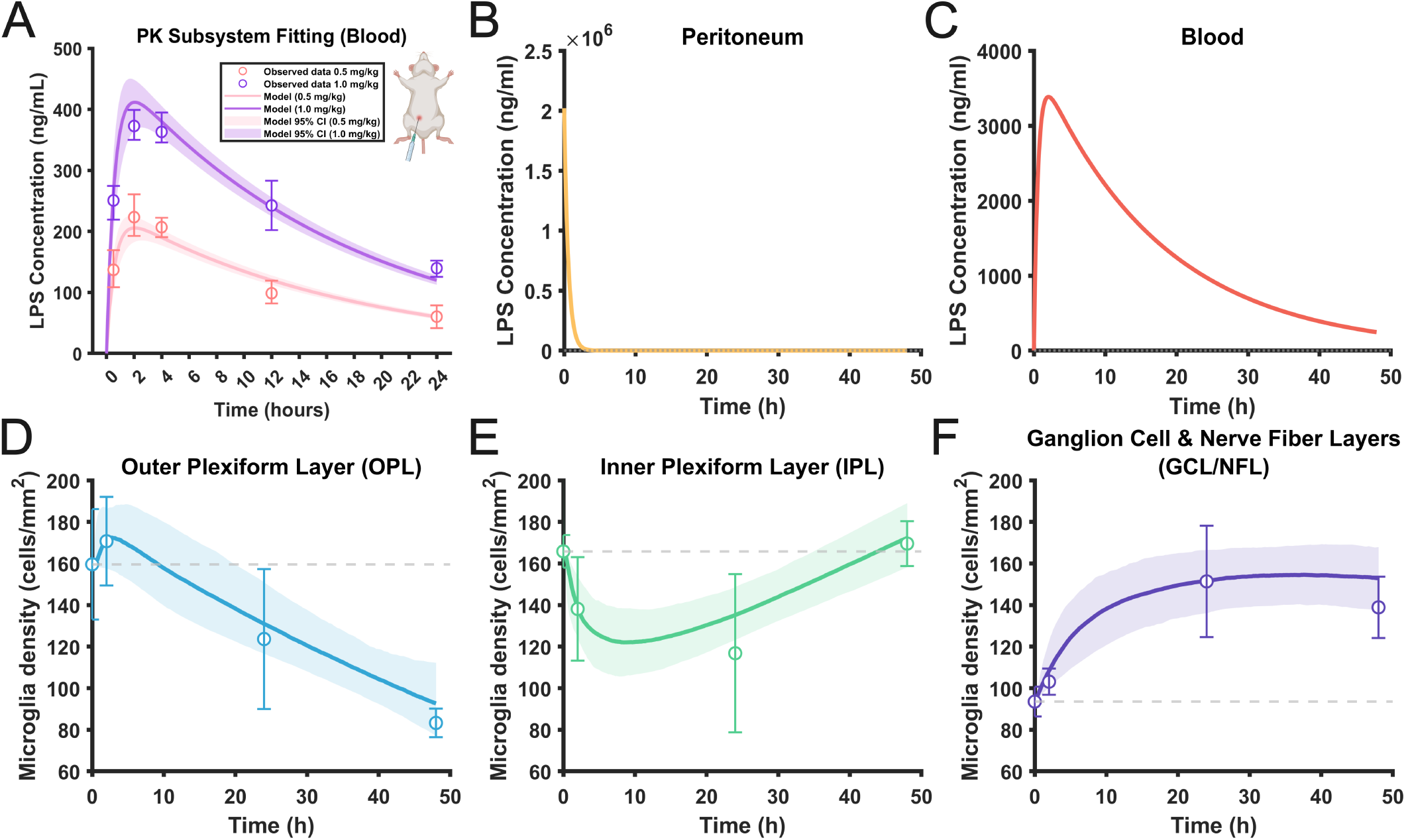
Model fits and posterior predictive bands following intraperitoneal LPS treatment. (**A**) Model fits of the PK system (**Equations 1–2**) to plasma LPS concentration data following intraperitoneal administration are shown for two doses (0.5 mg/kg and 1.0 mg/kg) reported by Huang *et al*. [42]. Shaded bands represent approximate 95% confidence intervals obtained using a nonlinear least-squares covariance approximation and propagated through the model. (**B–C**) Simulated LPS concentrations in the peritoneum and blood following 9 mg/kg IP administration. (**D–F**) Model fits to measurements reported by Dando *et al*. [15] for microglia densities in the outer plexiform layer (OPL), inner plexiform layer (IPL), and ganglion cell layer/nerve fiber layer (GCL/NFL) following intraperitoneal administration of 9.0 mg/kg LPS. Solid curves denote posterior mean model predictions, while shaded regions indicate Bayesian posterior predictive intervals (2.5th–97.5th percentiles) obtained by propagating posterior parameter samples through the coupled PK–microglia migration model. This figure is supported by the following supplementary information: **Table supplement 2: Fitting statistics for the microglia migration model following IP LPS administration**.

### 2.7 Rat Pharmacokinetics

To evaluate the generalizability of the model beyond mice, parameters from the pharmacokinetic (PK) subsystem were re-fit using published plasma LPS measurements from rats [44]. Rats are widely used in endotoxin studies [59–62] and share many immunological features with mice, yet exhibit known differences in LPS response and clearance.

Consistent with our mouse parameterization, *Vol*_*p*_ was taken to represent the resting peritoneal fluid volume rather than the total volumetric capacity of the peritoneal cavity. In rats, this resting volume has been directly measured as 3.07± 0.18 mL [63], and we therefore set *Vol*_*p*_ = 3.07 mL. As Hutter & Kim [44] did not specify the injection volume used to administer the 200 *µ*g/kg LPS dose, we assumed it to be negligible. Given rats are substantially larger than mice and the injected volume is small relative to the resting peritoneal fluid volume this seemed justified. Blood volume (*Vol*_*B*_) in rats is well established and can be taken as approximately 0.064 mL/g [64]. Using the mean body mass of 283 grams reported for the rats in the dataset by Hutter & Kim [44], the total blood volume was estimated as *Vol*_*B*_ = 18.112 mL.

The blood LPS clearance rate *k*_*B*_ was estimated from published pharmacokinetic data describing plasma endotoxin concentrations in rats following intraperitoneal administration [44]. In that study, circulating LPS was quantified using glucosamine residue concentrations as a proxy for plasma endotoxin over time and *k*_*B*_ was calculated from the approximately exponential decline observed over the 8–24 h interval giving a half-life of 7.43 h [44], i.e. *k*_*B*_ = 0.0933 h^−1^. Importantly, this value should be interpreted as an effective elimination rate for use in the model, rather than a true terminal half-life. Parameters *F* and *k*_*R*_ were then refitted to the rat data [44] using log-transformed concentrations and model outputs. Nonlinear least-squares fitting yielded parameter estimates of *F* = 0.0013 and *k*_*R*_ = 0.1264. The full set of rat pharmacokinetic parameters is summarized below in Table 4.

**Table 4.** Parameter values used to recalibrate the rat pharmacokinetic subsystem. Parameters include fitted absorption and bio-availability terms, literature-derived physiological volumes, and an effective blood clearance rate estimated from the plasma endotoxin measurements of Hutter and Kim [44].

| Parameter | Value | Reference | Units |
| --- | --- | --- | --- |
| $k_R$ | 0.1264 | Fitted to data from [44] | $\text{h}^{-1}$ |
| $F$ | 0.0013 | Fitted to data from [44] | - |
| $Vol_P$ | 3.07 | Zakaria & Rippe [65] | mL |
| $Vol_B$ | 18.112 | Diehl [64]; Hutter & Kim [44] | mL |
| $k_B$ | 0.0933 | Explicit calculation from [44] | $\text{h}^{-1}$ |

### 2.8 Rhesus Macaque Pharmacokinetics and Microglia Initial Distributions

To simulate microglia migration in response to LPS stimuli in rhesus macaques, we combined plasma measurements for LPS injected intravenously in macaques [45], with initial microglia distributions in the retina of macaques [18]. As LPS was administered intravenously, the intravenous delivery model (**Equation 3**) was used to model LPS PKs. The model was fitted to plasma LPS concentration data using nonlinear least-squares estimation. The resulting parameter estimates are provided in Table 5.

**Table 5.** Fitted parameter values for the rhesus macaque intravenous LPS bi-exponential decay model (**Equation 3**) obtained from nonlinear least-squares fitting to plasma LPS concentration data from Mathison *et al*. [45] following intravenous administration in rhesus macaques.

| Parameter | Interpretation | Value | Units |
| --- | --- | --- | --- |
| $A_1$ | Fast-phase amplitude | 0.7146 | - |
| $A_2$ | Slow-phase amplitude | 0.2711 | - |
| $k_1$ | Fast clearance rate | 0.3027 | $\text{min}^{-1}$ |
| $k_2$ | Slow clearance rate | 0.0019 | $\text{min}^{-1}$ |

Furthermore, rhesus macaques were assumed to have a representative body weight of 6.75 kg, corresponding to the midpoint of the body weight range reported in the experimental study from which pharmacokinetic data were extracted. At a dose of 9 mg/kg, this yields a total administered mass of 60.75 mg of LPS. Blood volume in rhesus macaques was set at 54 mL/kg of body weight [66], giving a total blood volume of 364.5 mL. The initial blood concentration was therefore estimated as

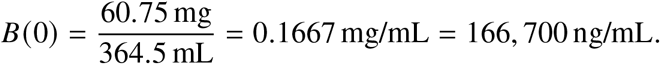

To mimic the retinal microscopy sampling procedure used by Dando *et al*. [15], in which microglia densities were quantified from randomly selected fields of view, our primary simulations used counts obtained from the peripheral retina (i.e., excluding the macula and fovea). This choice reflects the fact that the macula and fovea are classically defined as the central 5-6 mm radius region of the retina [67], corresponding to ≤ 10% of the total retinal surface area assuming a retinal area of approximately 1000–1100 mm^2^ [68]. Consequently, randomly sampled fields of view are most likely to capture peripheral retinal regions.

Singaravelu *et al*. [18] measured the initial density of microglia in the retinal layers in the fovea, macula and peripheral retina regions. We determined a kernel density approximation for their measurements using MATLAB’s ksdensity, from which we generated 1,000 samples for each retina layer and each retina region. These samples were used as initial conditions for the model.

## 3 Results

### 3.1 Model of Microglia Migration in response to LPS

To investigate retinal microglia migration following systemic LPS administration, we fit parameters in the IP injection and microglia migration subsystems to measurements in mice. Plasma LPS measurements reported by Huang *et al*. [42], were used to calibrate the PK subsystem (**Equations 1 -2**; Figure 3A). The migration subsystem (**Equations 4-6**) was then fit to experimental measurements from Dando *et al*. [15] (Figure 3B–F). Both subsystems accurately captured the experimental measurements. Each retinal layer exhibits a qualitatively different temporal response, and the model successfully reproduces all three behaviors. Furthermore, Figure 3B–C demonstrates that the predicted LPS trajectories align well with the timing of microglia redistribution hallmarks, such as the rapid initial drop in IPL microglia density, suggesting that the initial redistribution of microglia is tightly coupled to the concentration of circulating LPS. Quantitative fit statistics further support the qualitative goodness-of-fit observations (Table S3). RMSE values, representing the typical prediction error, remained small relative to microglia densities (6.5–8.5 cells/mm^2^), indicating high accuracy across all retinal compartments.

Predictive uncertainty in the microglia migration model was quantified using Bayesian posterior predictive simulations (Figure 3D–F). The resulting 2.5th–97.5th percentile predictive intervals capture the general variability observed across retinal layers, though several individual animal measurements fall outside the predicted ranges. This suggests that while the calibrated parameter ranges adequately describe typical responses, they do not fully capture the more extreme responses observed in some animals. This observation motivated the extended uncertainty quantification presented in Figure S5, where parameter ranges were deliberately widened to explore less common individual responses.

### 3.2 Effects of LPS Dose and Bolus Schedule on Retinal Microglia Migration

Following validation of the mathematical model, we next examined how microglia migration dynamics change with LPS dose size and schedule. We first simulated the model under eight escalating doses and a no-LPS control (Figure 4A). Decreasing the endotoxin dose from 9.0 mg/kg produced a stronger preference for microglia migration from the OPL to the IPL. This behavior arises from the dose-dependent structure of the model’s migration fluxes. At low doses (lighter curves in Figure 4A), increasing endotoxin levels strengthens microglia migration from the OPL to the IPL as the activation governing OPL efflux exceeds the corresponding influx term, reflecting the asymmetry *η*_*O*_ *≫ η*_*G*_ (Figure S4). However, as the dose becomes large, both activation terms approach saturation and their influence weakens. In this regime, the intrinsic migration rates dominate, with OPL influx (*k*_*IO*_) approaching OPL efflux (*k*_*OI*_ ). This shifts the net migration balance back toward the OPL, producing the observed reversal and the non-monotonic dose–response relationship in OPL microglia density predicted by the model.

**Figure 4:**
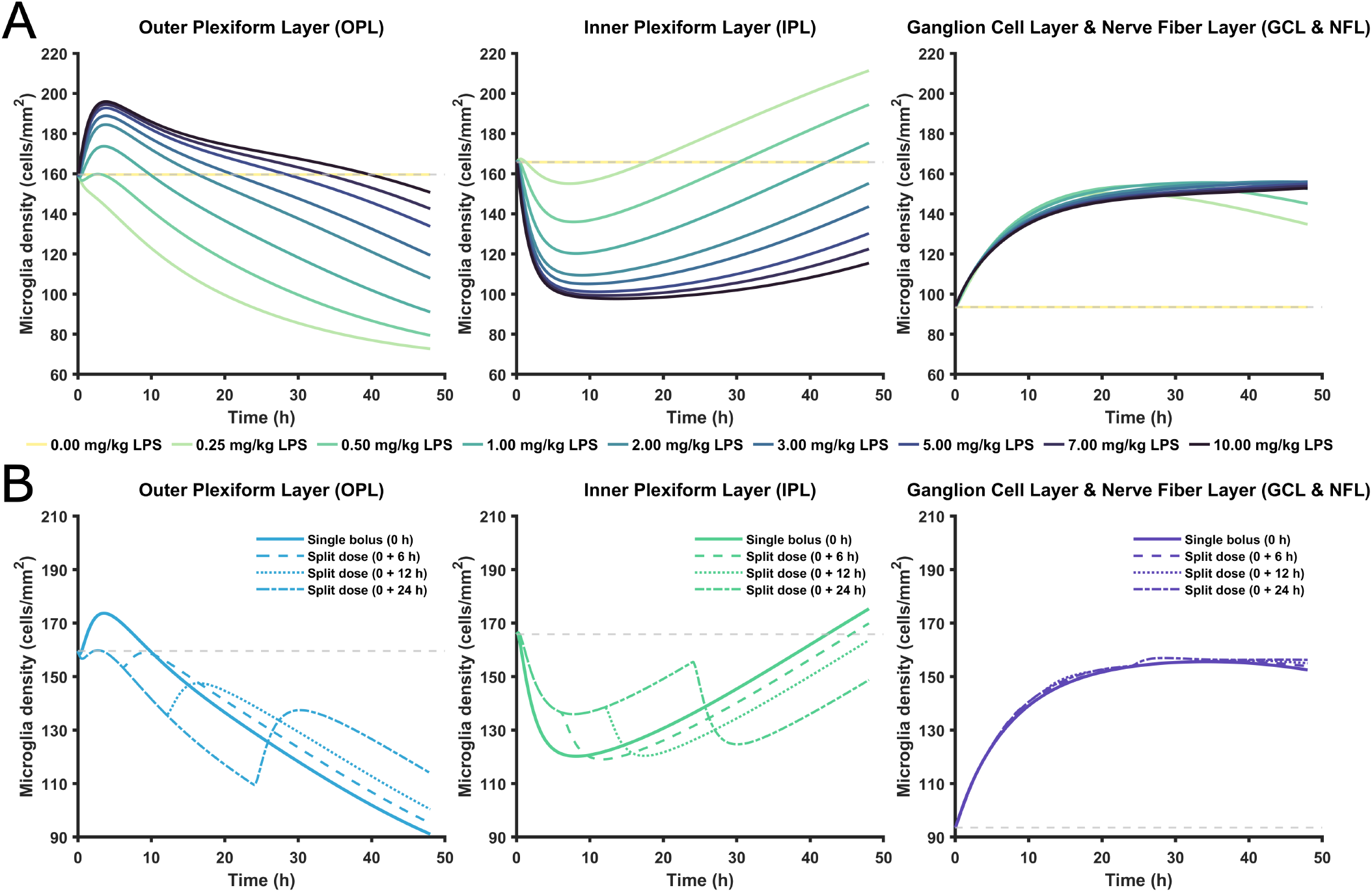
Parameter sweeps of intraperitoneal LPS dose level and administration schedule. (**A**) Model simulations depicting the response of the microglia migration subsystem to changes in the initial peritoneal LPS dose across an experimentally relevant range. Trajectories are shown for the OPL, IPL, and GCL/NFL, with each sub-panel corresponding to one retinal compartment. **(B)** Comparison of intraperitoneal dosing schedules demonstrating the effect of administering a total 9 mg/kg LPS dose as either a single bolus or as multiple smaller injections delivered at later time points (0 & 6 h, 0 & 12 h, or 0 & 24 h) Panels show the resulting microglia trajectories and the underlying peritoneal and blood LPS profiles for each dosing schedule. This figure is supported by the following supplementary information: **Figure supplement 4: Additional model variables during intraperitoneal LPS dose escalation**.

To evaluate how repeated LPS exposure influences predicted microglia redistribution, sequential LPS administrations were simulated by increasing peritoneal LPS concentrations at specified time points and solving the full model in a piecewise manner (Figure 4B). Experimental evidence suggests that endotoxin tolerance may begin to emerge as early as approximately 1h following LPS treatment, with attenuated inflammatory responses observed at later times [69, 70]. Nevertheless, simulating repeated LPS treatment remains informative for exploring how successive stimuli may interact in the absence of explicit LPS tolerance. Sequential dosing events following the initial dosage were implemented at 6h, 12h, and 24h. The resulting microglia density dynamics displayed a consistent pattern across layers. In the OPL, all split-dose regimens produced a more prolonged decline than the single bolus. Conversely, the IPL showed more sustained depletion when the second dose was administered early.

In contrast to both plexiform layers though, the GCL/NFL trajectories were strikingly insensitive to dosing schedule. Furthermore, the GCL/NFL saturates heavily at higher LPS doses (Figure 4A), reflecting a broader resilience of GCL microglia density to changes in dosage. Mathematically, GCL/NFL saturation occurs because the Hill activation terms saturate at high LPS concentrations, so further increases in *B*_0_ no longer increase migration into *G*, leaving the dynamics governed by the intrinsic kinetic rates and logistic stabilization and causing *G*(*t*) to plateau.

### 3.3 Parameter Influence on Microglia Migration

We next wanted to assess which biological pathways, represented by parameters in our model, were most responsible for the density of microglia in each layer. To do this, we conducted a sensitivity analysis using partial rank correlation coefficients (PRCCs) (Figure 5). This analysis revealed that parameters governing intrinsic microglia migration exert a substantially greater influence on retinal microglia density than parameters describing LPS clearance and systemic distribution. In particular, increases in LPS-related parameters beyond the initial inflammatory signal produced relatively modest changes in microglia outcomes compared to perturbations in migration rates. Together, these results indicate that once inflammation is initiated, the ensuing redistribution of microglia is driven primarily by intrinsic, retina-local migration dynamics rather than by additional increases in LPS exposure. While this suggests a dominant role for intrinsic migration dynamics, it also raises the possibility that the weak sensitivity to LPS-related parameters may reflect saturation of the Hill activation terms. To assess this, we further examined sensitivity to average and peak values of 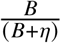 . These quantities exhibited near-zero PRCC values with respect to GCL/NFL AUC, suggesting that the model indeed operates in a regime where LPS-dependent migration is highly saturated, as expected under the large dose of LPS (9 mg/kg) used in these simulations.

**Figure 5:**
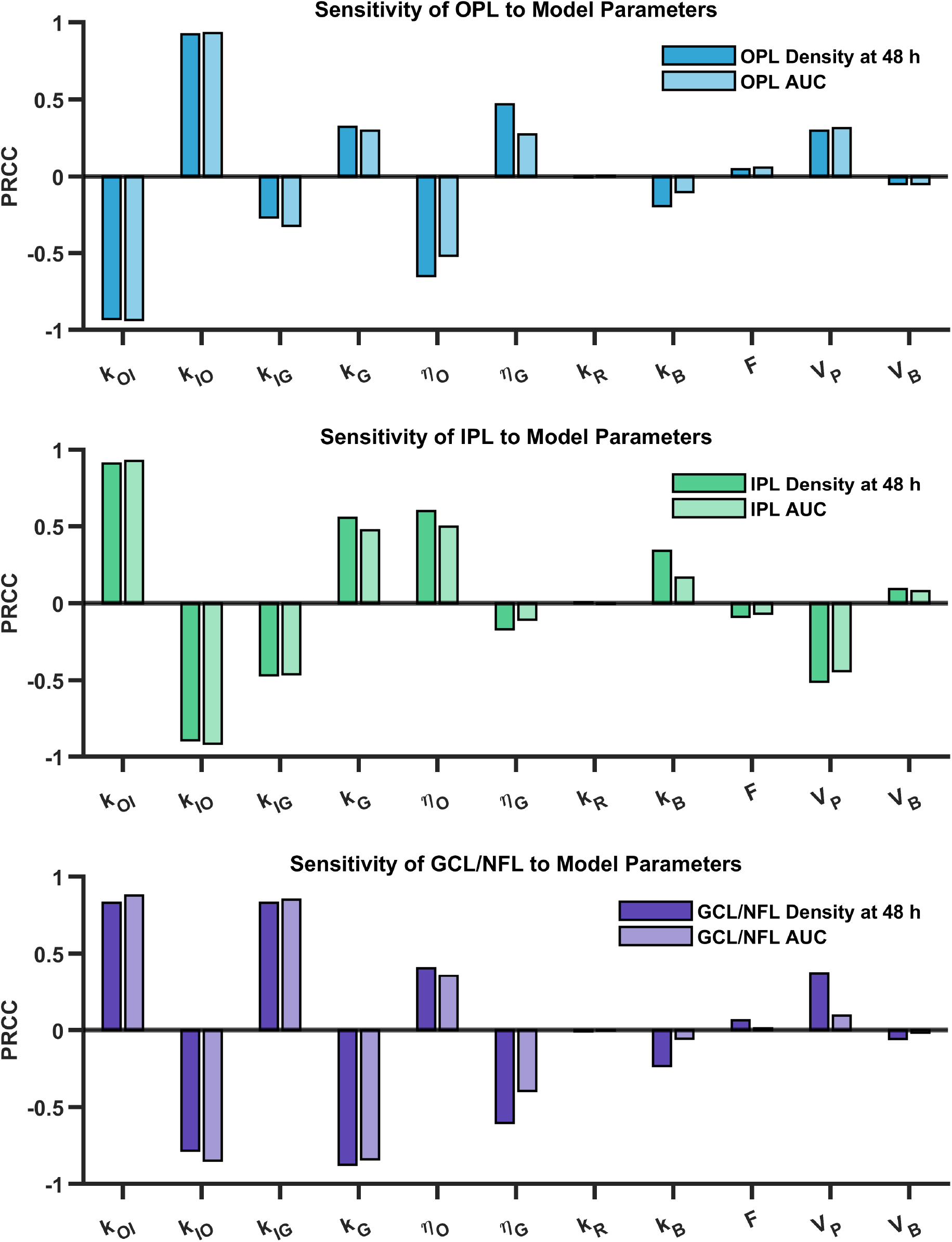
Global sensitivity analysis of parameters in the coupled PK–microglia migration model following IP LPS administration. Partial rank correlation coefficients (PRCCs) quantifying how variations in model parameters influence microglia density outcomes in each retinal layer. Sensitivity of OPL (top), IPL (middle) and GCL/NFL (bottom) microglia density outcomes, shown for final density at 48 h and the 0–48 h area under the curve (AUC). Positive coefficients indicate parameters that increase the outcome measurement when increased within their sampling range, whereas negative coefficients indicate inverse relationships. Analyses were performed using Latin hypercube sampling over the parameter ranges listed in Table 3, with 100,000 model simulations used to compute PRCCs. This figure is supported by the following supplementary information: **Figure supplement 5: Extended global sensitivity analysis using widened parameter ranges.** **Figure supplement 6: Monotonicity assessment for partial rank correlation coefficient analysis**.

A further insight from the PRCC analysis is that microglia exchange between the OPL and IPL exerts a substantially stronger influence on model outcomes than exchange between the IPL and GCL/NFL. Parameters governing bidirectional migration between the OPL and IPL (*k*_*OI*_ and *k*_*IO*_) exhibit the largest absolute PRCC values across both 48 h density and AUC metrics in these layers, indicating that redistribution within the plexiform layers dominates the control of microglia density. In contrast, parameters controlling IPL–GCL/NFL exchange (*k*_*IG*_ and *k*_*GI*_ ) consistently show weaker correlations with microglia density outside of the GCL/NFL, suggesting that fluxes into and out of the GCL/NFL play a secondary role in shaping microglia density trajectories outside of this layer.

Finally, extending the sensitivity analysis to substantially widened parameter ranges did not largely alter these conclusions (Figure S5). However, the extended analysis did reveal an increased sensitivity of GCL/NFL microglia density to PK parameters, suggesting that extreme changes to the concentration of circulating LPS are needed to break down this compartment’s resilience to dose scaling. The validity of the PRCC analyses (Figure 5; Figure S5) depends on the monotonicity of model outputs as parameters vary. This requirement was verified using monotonicity plots (Figure S6), which are provided in the interest of methodological rigor.

### 3.4 Microglia Redistribution Depends on LPS Delivery Route

Although dose magnitude is the most readily adjustable aspect of LPS administration, the route of delivery is also expected to substantially influence inflammatory dynamics. To examine this, we fitted the PK model for IV administration (**Equation 3**), formulated as a biphasic decay model (see *Methods*), to plasma LPS measurements in mice following a 9.0 mg/kg injection [15] and compared the resulting microglia responses to those produced by the same dose delivered intraperitoneally (Figure 6A). The IV model demonstrated excellent qualitative and quantitative agreement with experimental data, validating its use for simulating intravenous-LPS blood concentration.

**Figure 6:**
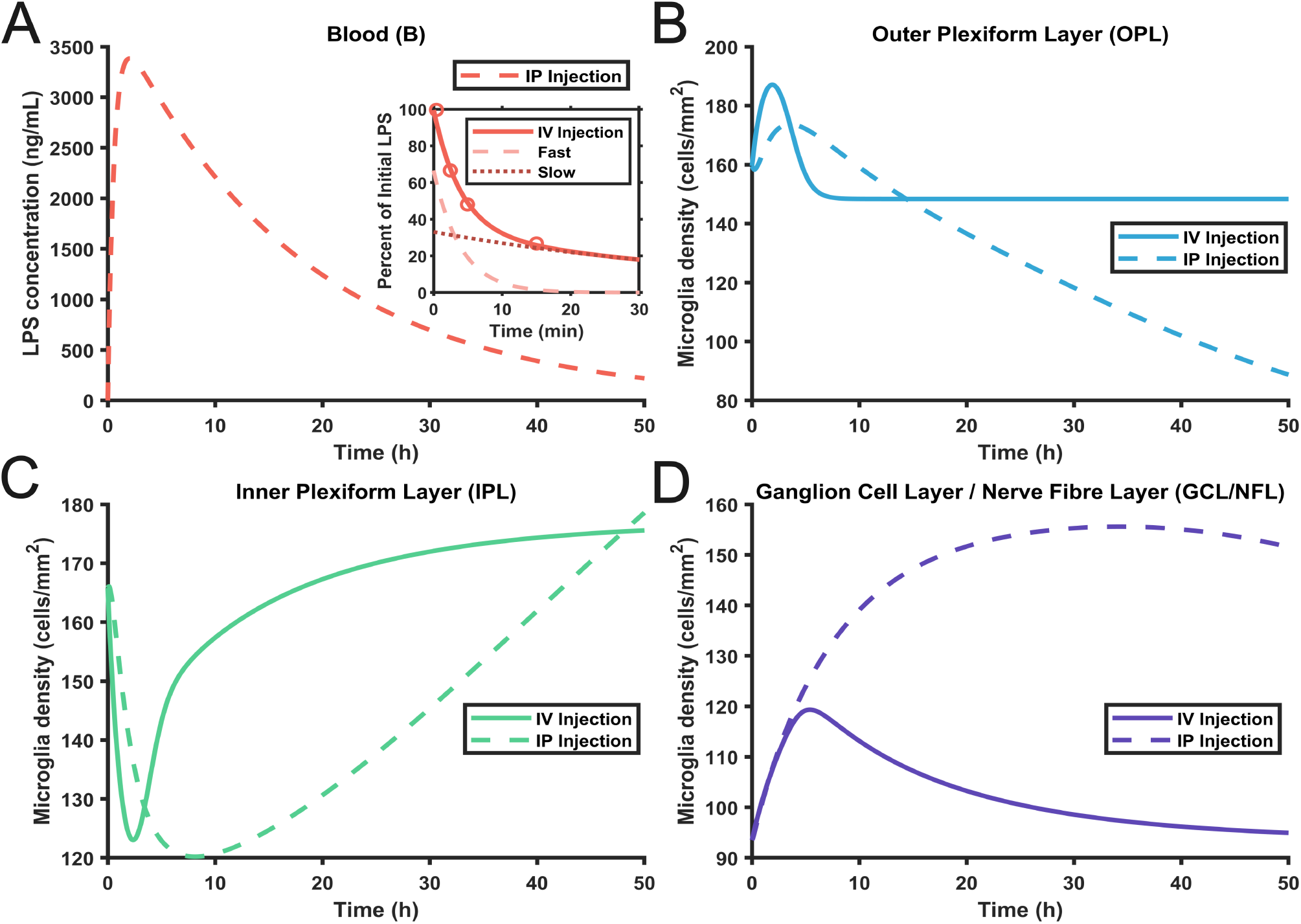
Comparison of the effects of intraperitoneal versus intravenous LPS administration on retinal microglia redistribution in mice. **(A)** The IV subsystem (**Equations 3**) was fit to blood LPS concentrations following intravenous injection [43]. The fast and slow decay components are shown separately (dashed & dotted curves) along with their combined fit (solid curve) in the figure insert. The fitted IP subsystem from Figure 3 is shown as a dashed line in the main figure. **(B-D)** Comparison of the effect of intravenous and intraperitoneal LPS administration on microglia migration dynamics under matched 9 mg/kg treatment. Solid curves show intravenous LPS–driven microglia densities, while dashed curves show the corresponding intraperitoneal LPS-driven densities.

Despite receiving the same total dose, the IV injection induced a brief, high-amplitude spike in blood LPS concentration, whereas IP administration generated a more delayed and prolonged profile, consistent with the characteristic pharmacokinetics of each route. These contrasting profiles led to highly different time courses of microglia migration. In the OPL, IV delivery elicited a rapid overshoot followed by rapid resolution, while IP delivery drove a monotonic decline throughout the entire 48h window. The IPL exhibited a contrasting behavior where IV dosing caused a rapid early depletion with gradual recovery toward baseline, whereas IP induced these changes in a smoother and more gradual fashion. In the GCL/NFL, IV injection produced only a small and transient spike in microglia density, whereas IP injection resulted in a much larger and sustained accumulation of microglia. These results suggest that microglia redistribution is more strongly shaped by the duration of inflammatory exposure and hence dependent on LPS delivery route.

### 3.5 Predicted Variability Between Mouse and Rat Responses to Intraperitoneal LPS

Thus far, the effects of LPS dose, biological pathways, administration route, and bolus timing on microglia migration trajectories have all been examined. A natural next question is how microglia migration generalizes across species, particularly given known differences in species LPS pharmacokinetics. To address this, we refit the IP PK subsystem of our model using published rat plasma LPS data (Figure 7A; [44]) and simulated the model under rat-specific pharmacokinetic parameters (Figure 7B-F; Table 4).

**Figure 7:**
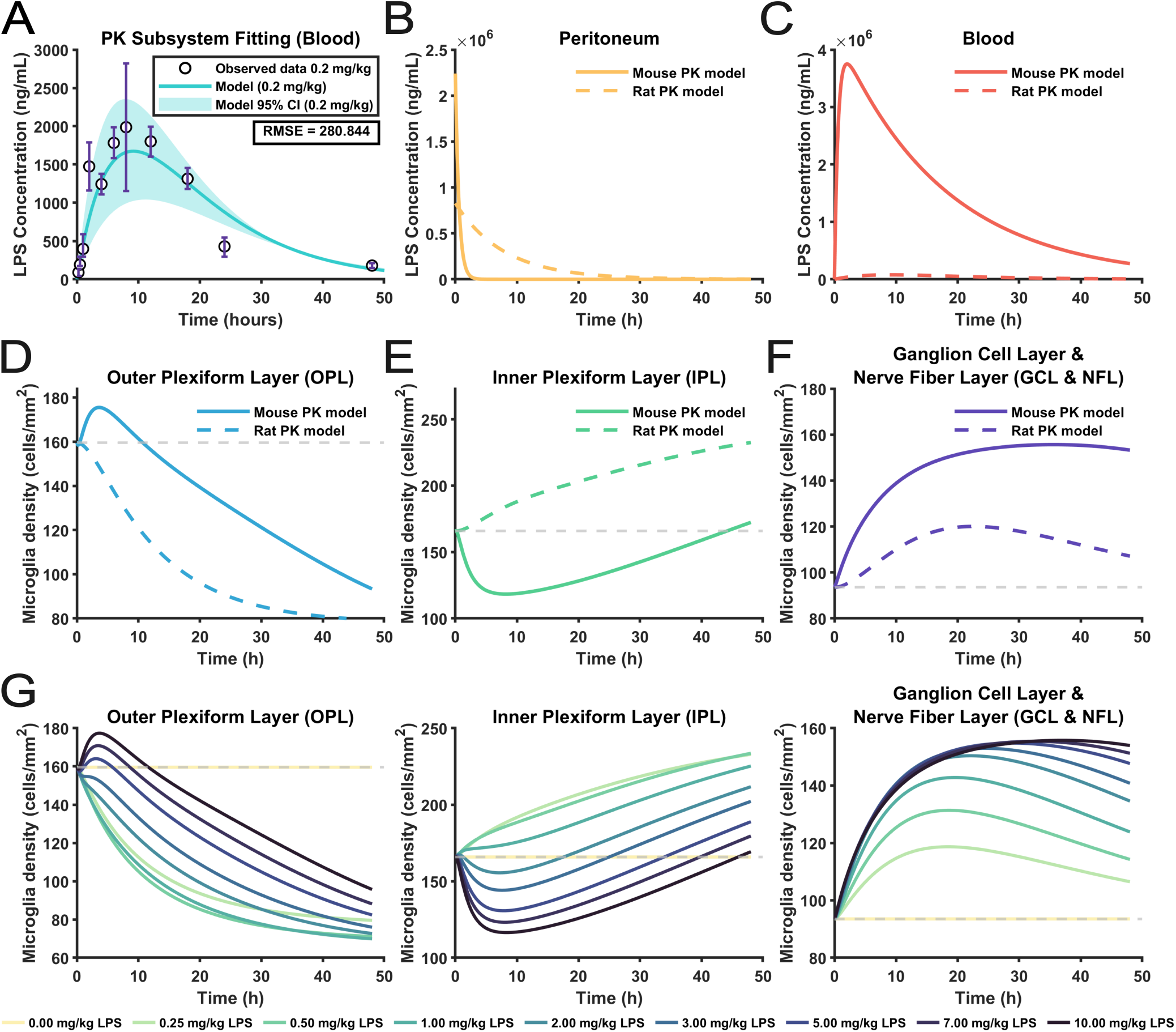
Cross-species model comparison between mice and rats. **(A)** Recalibration of the PK subsystem to rats injected intraperitoneally with LPS. The best-fit plasma LPS curve to blood LPS measurements [44] is shown together with approximate 95% uncertainty. **(B)-(F)** Cross-species comparison between the rat model (bold curves) and the previously parameterized mouse model (light curves) following a dose of 9.0 mg/kg. **(G)** Model simulations depicting the response of the microglia to changes in the initial intraperitoneal LPS dose (0–10 mg/kg) in rats. All simulations were performed under the parameter regime provided in Table 4.

The resulting simulations revealed several differences in predicted microglia behavior between rats and mice, driven entirely by the distinct PK profiles of the two species (Figure 7B-F). Rats exhibit lower LPS bioavailability, slower peritoneal absorption, and faster blood clearance, systemic LPS exposure is both weaker and shorter-lived in the rat-fitted model than in mice. This shortened duration of inflammatory stimulus produced correspondingly shorter migration responses across all layers. In the OPL, rats showed a faster decline in microglia density, quickly approaching an equilibrium far below baseline. The IPL in rats exhibited slowly increasing density due to influx from the other layers, in contrast to the drop and rebound that is predicted in mice. Similarly to the OPL, the accumulation of microglia in the GCL/NFL was attenuated in rats, with a lower peak magnitude and quicker return toward more baseline conditions. More broadly, these predictions suggest that species-specific pharmacokinetics alone can reshape the trajectory of microglia migration responses, even when the underlying migration dynamics are held constant.

To contextualize how species-specific differences relate to dose sensitivity, we next simulated a wide range of intraperitoneal LPS doses using the rat-fitted PK–migration system (Figure 7G). Increasing the administered dose produced the expected monotonic elevation in both peritoneal and blood LPS concentrations. These dose-dependent PK profiles translated into mostly monotonic changes in microglia redistribution across all retinal layers, with the magnitude of peaks being proportional to circulating LPS concentrations. The IPL, however, exhibited a distinct nonmonotonic behavior as doses increased. At low escalating doses, IPL microglia density rose monotonically, as progressively larger doses below approximately 3 mg/kg caused more migration into this layer from the OPL. At higher doses however, IPL microglia density curves began to exhibit a brief dip between about 4h and 12h post treatment due to the much higher efflux from the IPL into the neighboring GCL.

### 3.6 Recalibration to Rhesus Macaque and Sensitivity to Initial Microglia Distributions

Model behavior was previously examined under fixed baseline microglia distributions in rodents; however, measurements for the density of microglia across the layers are available in macaques [18] (Figure 8A-B). To use these measurements to assess the impact of the initial microglia distribution on migration dynamics following IV injection, we first re-fit the IV subsystem to plasma LPS measurements in macaques [45].

**Figure 8:**
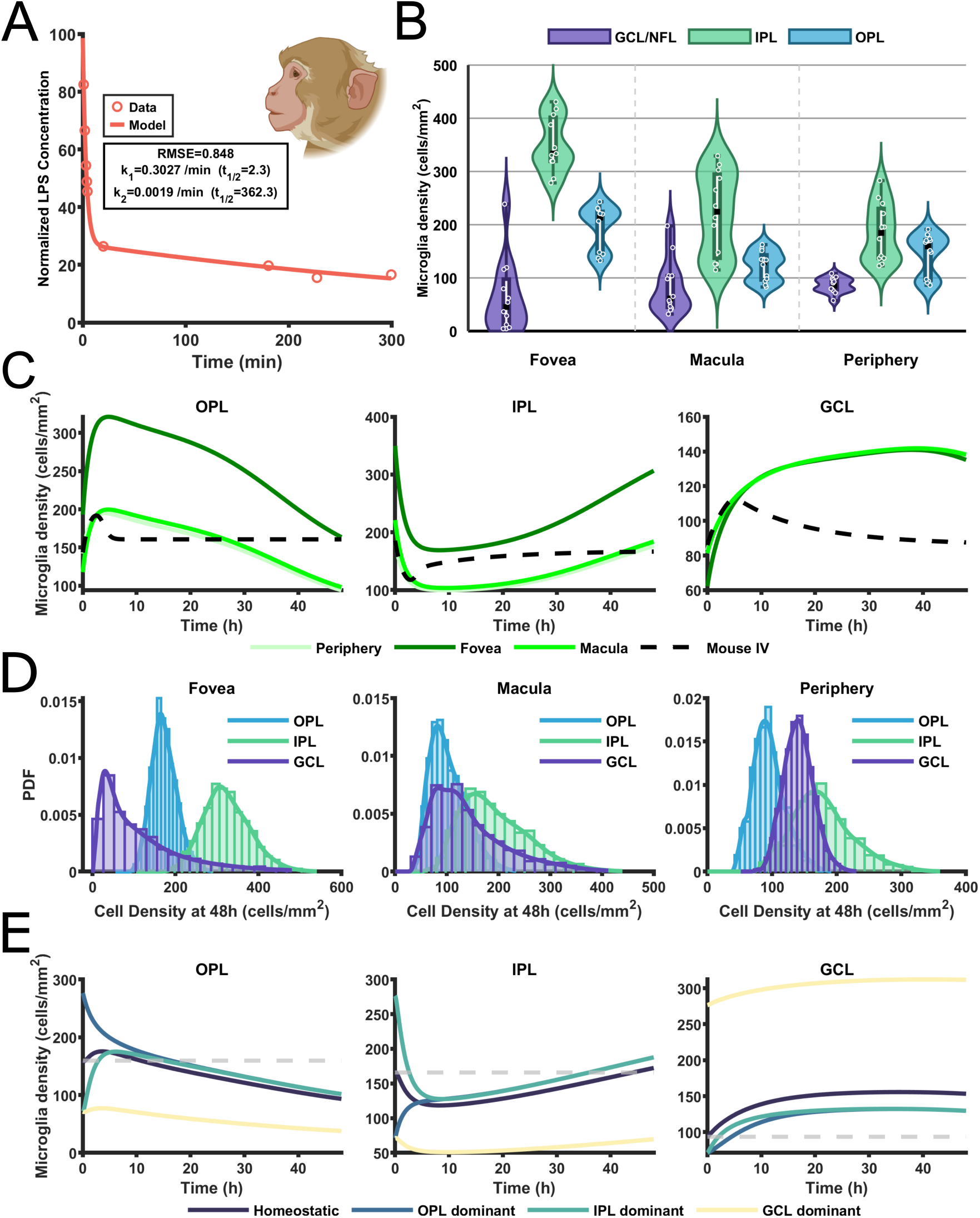
Influence of initial microglia distributions on retinal migration dynamics. (**A**) Biphasic decay model fitted to experimentally measured blood LPS concentrations following intravenous injection in rhesus monkeys [45]). (**B**) Microglia density measurements in untreated rhesus macaque retina [18], shown as violin plots with embedded box plots and individual data points for each retinal region (fovea, macula, and periphery). Densities are reported separately for the GCL/NFL, IPL, and OPL layers. (**C**) Model simulations of retinal microglia dynamics using the average region-specific initial conditions from (**B**). Trajectories for foveal, macular, and peripheral initial conditions are compared with the original mouse IV simulation for reference. (**D**) Distributions of model-predicted microglia densities at 48 h obtained by sampling initial conditions from kernel density estimates (KDEs) of the data shown in **B**. Separate distributions are shown for the OPL, IPL, and GCL/NFL layers across the three retinal regions. (**E**) Simulations of the model for the experimentally observed initial condition, OPL-dominant, IPL-dominant, and GCL-dominant configurations. This figure is supported by the following supplementary information: **Figure supplement 7: Prolonged microglia migration in rhesus monkeys arises from sustained blood LPS despite lower early-time concentrations**. **Figure supplement 8: Propagation through the model smooths heterogeneous initial microglia distributions into unimodal outcomes**.

Using the macaque-calibrated intravenous model, microglia migration dynamics were compared between mice and rhesus macaque under a matched intravenous LPS treatment of 9 mg/kg (Figure 8C) for average initial densities taken from the periphery, fovea and macula. Despite identical delivery routes, the two species exhibited different migration trajectories driven by their respective blood LPS kinetics.Macaques consistently exhibited more sustained and gradual redistribution dynamics, whereas mice showed transient peaks followed by faster recovery. These differences indicate that species-specific pharmacokinetics alone can reshape microglia redistribution patterns despite the same migration kinetics and LPS delivery route. Seeding the simulations with foveal, macular, and peripheral initial densities produced quantitatively different but qualitatively similar trajectories across all layers (Figure 8C). Variations in baseline densities almost exclusively shifted the magnitude of microglia redistribution without substantially altering other features.

To further investigate the impact of the initial microglia density in each retinal layer, we generated 1,000 samples from a kernel density estimate (KDE) of the microglia density measurements in each retinal layer [18] (Figure 8B). We then simulated the model for each of the 1,000 samples to obtain a distribution for microglia in each retinal layer after 48 hours (Figure 8D). Despite the clearly bimodal initial distribution of microglia in the data, the model yielded relatively narrow and unimodal distributions of predicted microglia densities at 48 h (Figure 8D). Comparing the foveal, macula and periphery regions, we see the macula and periphery regions had similar distributions of microglia across the retinal layers at 48 hours, however, the fovea was qualitatively difference, with low numbers of microglia in the GCL/NFL and high numbers in the IPL.

Finally, we considered the impact of extreme initial microglia distributions. We fixed the total microglia population *M*_*tot*_ and redistributed microglia to be dominant in one layer at a time. For example, in the OPL dominant scenario, the initial density of microglia in the OPL was 2 /3 ×*M*_*tot*_, and 1/ 6× *M*_*tot*_ in the remaining layers. This same redistribution was repeated for the IPL dominant and GCL dominant scenarios (Figure 8E). Across the three scenarios and the experimentally observed scenario, the model converged toward similar qualitative migration patterns, indicating a resilience of redistribution dynamics to initial layer density. The lone exception occurred when microglia were initially concentrated in the GCL/NFL, which produced a persistently GCL-dominant trajectory throughout the inflammatory response. This suggests that only extreme bias toward the GCL/NFL can alter qualitative redistribution outcomes.

## 4 Discussion

In this study, a mechanistic mathematical model was developed to describe how microglia redistribute across layers of the retina in response to inflammation. By coupling a pharmacokinetic model of LPS levels to a migration model describing inter-layer microglia movement, the model successfully recapitulated experimental data both qualitatively and quantitatively across all retinal layers. The model was able to capture both intraperitoneal and intravenous injection plasma measurements in mice, rats and rhesus macaques. The simulations then revealed how the dose, the route of administration, and species-specific pharmacokinetics alter the exposure to retinal LPS and subsequent microglia redistribution.

Sensitivity analyses demonstrated that intrinsic microglia migration parameters exert a substantially stronger influence on retinal microglia density than parameters governing LPS pharmacokinetics. While LPS dynamics are necessary to initiate microglia redistribution, variations in migration rates produced larger and more consistent effects on both 48h densities and cumulative activation than comparable changes in LPS availability parameters. A natural explanation for this behavior is that migration quickly saturates with higher LPS levels. This interpretation is consistent with experimental observations that exceptionally low endotoxin doses (as low as 0.05 mg/kg) are sufficient to elicit robust microglia reactivity [71]. At commonly used doses, further increases in LPS produce diminishing effects, making intrinsic migration kinetics the primary drivers of retinal microglia redistribution during inflammation.

Considering well-established species-specific differences in LPS response [72, 73], together with the fundamentally different goals of intraperitoneal versus intravenous delivery, it is not surprising that distinct microglia density trajectories emerge across delivery routes and experimental species. Intravenous administration is typically chosen to produce rapid systemic exposure, whereas intraperitoneal delivery yields a more delayed and prolonged inflammatory profile, and similar divergence is expected across species with different clearance and distribution kinetics. What is notable, however, is the extent to which these differences translate into qualitatively distinct microglia redistribution profiles. Our results demonstrate that modest changes in pharmacokinetic structure alone are sufficient to generate pronounced differences in the timing, persistence, and layer-specific organization of microglia responses, even when the underlying migratory parameters are conserved. These findings illustrate that the delivery route and choice of experimental species do not just rescale inflammatory responses, as is sometimes expected in pharmacology, but fundamentally reshapes retinal microglia redistribution.

A final yet pervasive observation from this study was the limited influence of baseline microglia layer density on the microglia migration. Across a wide range of initial layer densities, including those experimentally derived from rhesus monkeys and artificially generated to reflect extreme scenarios, the mathematical models consistently demonstrated a tendency to converge toward a common redistribution trajectory. From an experimental perspective, this relative insensitivity to baseline distributions has important implications. It suggests that inter-individual variability in resting microglia densities, potentially arising from age, sampling location, or unknown factors, may exert relatively little influence on the qualitative and quantitative features of microglia redistribution. The observed robustness to initial conditions further suggests a high likelihood of reproducibility in studies on microglia redistribution tendencies, even across cohorts with substantial heterogeneity.

### Limitations and Future Work

Several limitations of this study should be acknowledged. Across all mathematical models and parameterizations presented in this article, microglia migration kinetics and their associated parameters were assumed to be identical, reflecting the limited availability of data to support more specific or context-dependent parameterizations. Furthermore, the experimental data used for both the initial mouse model calibration and the rhesus monkey dataset relied on Iba-1 immunostaining, which is not specific to microglia during inflammation and may also label other myeloid cells present in small numbers within the CNS, including monocytes and neutrophils [74]. In addition, the model neglects microglia proliferation, death, and migration to compartments outside the modeled retinal layers, such as the subretinal space. Finally, because this study relies on the use of mathematical models as representations of biological systems, the results should be interpreted largely as a predictive line of evidence rather than definitive descriptions of microglia behavior.

Future work could extend this framework by integrating microglia phenotypes and coupling migration dynamics to transcriptional or phenotypic switching. Experimentally, longitudinal measurements of microglia fluxes between layers, as well as perturbations targeting intrinsic migration pathways, would provide valuable data to assess whether the model’s predictions would hold. Ultimately, combining our demonstrated mathematical techniques with fluorescent imaging and single-cell transcriptomics would enable future models to directly link retinal layer redistribution dynamics to functional microglia states, in line with broader field efforts [75–77].

Prior mathematical models have addressed related aspects of inflammation-driven immune dynamics in the eye and central nervous system. For example, Nicholson *et al*. modeled leukocyte trafficking into the eye during experimental autoimmune uveitis and explicitly incorporated the role of the blood-retina barrier in peripheral immune infiltration [78]. While conceptually adjacent, that framework focuses on immune cell entry rather than redistribution of resident microglia. In contrast, the present modeling framework assumes that no peripheral immune cells infiltrate the OPL, IPL, GCL, or NFL during acute inflammation and that no new microglia proliferate within the first 48h following LPS exposure. A natural extension of our work could be to combine the model from Nicholson *et al*. with ours to capture the effects of inflammatory stimuli on microglia over a longer time frame.

## Acknowledgements

The authors acknowledge the support of QUT eResearch and Central Analytic Research Facilities (CARF).

A.J and T.J. also acknowledge research support from the Australian Research Council (ARC) #DE240100650. TC is grateful for partial support from the QJMAM Fund for Applied Mathematics and the London Mathematical Society.

## Author Contributions

T.J., T.C., and A.J. developed the mathematical model, conducted simulations, and interpreted analyses. S.D. contributed original data and biological insight. T.J., T.C., S.D., and A.J. provided substantial contributions to the discussion of the article’s content. T.J. drafted the article. S.D., T.C., and A.J. reviewed and edited the manuscript prior to submission. All authors substantively contributed to research design.

