## Supplementary Information for "Linking Systemic Endotoxin Exposure to Retinal Microglia Migration Through Mathematical Modeling"

### Mathematical Modeling of Retinal Microglia in Response to Endotoxin-induced Inflammation

This supplementary document provides methodological details and supporting results that complement the main article. Specifically, in Section S1, we solve the intraperitoneal PK model provided by **Equations 1-2** from the main text and show how it relates to the bi-exponential model describing intravenous injections. In Section S2, we provide details of how we calculated the model initial conditions. In Section S3, we present a list of alternative ODE models that were tested against the microglia density measurements from Dando *et al.* [1]. We emphasize that Section S3 does not represent a rigorous model selection process, but instead provides more detail surrounding how we arrived at the final model presented in the main text. In Section S4, we discuss the justification for our choice of $\eta_O$.

In Section S5, we present additional figures and tables supporting the methods and results of the main text. We include a table summarizing fitting statistics for the mathematical models describing microglia migration following LPS treatment. We conduct a parameter sweep that aids in the interpretation of results presented in **Figure 5** of the main text. We also provide an extended sensitivity analysis in which PRCCs were computed using parameter ranges that reflect biologically extreme or uncommon scenarios. Supporting plots are also included to confirm that the monotonicity assumption for valid PRCC interpretation is satisfied. To further explain the species-specific differences in microglia dynamics described in the main text, we present blood LPS concentration profiles comparing the mouse and monkey PK models across two time windows. Finally, we present ensemble model trajectories simulated from kernel density estimate (KDE)-sampled initial microglia distributions, demonstrating that model propagation smooths heterogeneous initial conditions into unimodal outcomes within each retinal layer.

#### S1 Connecting the Intraperitoneal PK model to the Biexponential PK Model

The biexponential form used to represent intravenous LPS pharmacokinetics (**Equation 3**) can be interpreted as a phenomenological expression consistent with the dynamics of the mechanistic two-compartment pharmacokinetic model (**Equations 1–2**). This relationship can be seen directly by solving the intraperitoneal pharmacokinetic subsystem:

$$\frac{dR}{dt} = -k_R R, \tag{S1}$$

$$\frac{dB}{dt} = F k_R \left(\frac{Vol_P}{Vol_B}\right) R - k_B B. \tag{S2}$$

**Equation S1** has the solution

$$R(t) = R(0)e^{-k_R t}. \tag{S3}$$

Substituting this into the blood compartment **Equation S2** and rearranging gives

$$\frac{dB}{dt} + k_B B = F k_R \left(\frac{Vol_P}{Vol_B}\right) R(0) e^{-k_R t}.$$

Multiplying through by the integrating factor $e^{k_B t}$ gives

$$e^{k_B t}\frac{dB}{dt} + k_B e^{k_B t} B = F k_R \left(\frac{Vol_P}{Vol_B}\right) R(0) e^{(k_B - k_R)t},$$

which can be written as

$$\frac{d}{dt}\left(B e^{k_B t}\right) = F k_R \left(\frac{Vol_P}{Vol_B}\right) R(0) e^{(k_B - k_R)t}.$$

Integrating both sides gives

$$B e^{k_B t} = \frac{F k_R \left(\frac{Vol_P}{Vol_B}\right) R(0)}{k_B - k_R} e^{(k_B - k_R)t} + C,$$

where $C$ is a constant of integration. Dividing through by $e^{k_B t}$ gives

$$B(t) = \frac{F k_R \left(\frac{Vol_P}{Vol_B}\right) R(0)}{k_B - k_R} e^{-k_R t} + C e^{-k_B t}.$$

Applying the initial condition $B(0)$ gives

$$C = B(0) - \frac{F k_R \left(\frac{Vol_P}{Vol_B}\right) R(0)}{k_B - k_R},$$

so

$$B(t) = B(0) e^{-k_B t} + \frac{F k_R \left(\frac{Vol_P}{Vol_B}\right) R(0)}{k_B - k_R} \left(e^{-k_R t} - e^{-k_B t}\right). \tag{S4}$$

Thus, the mechanistic two-compartment model in **Equations 1-2** naturally produces a biexponential blood concentration profile, i.e. **Equation S4**. The intravenous model (**Equation 3**) adopts this same biphasic functional form phenomenologically, without explicitly modeling the underlying compartmental transfer process. Note, while the general form of **Equation S4** and **Equation 3** are equivalent, the models capture different pharmacokinetic assumptions, i.e., IP injection PKs are dominated by transfer from the peritoneal to the blood and a single clearance rate, whereas IV injection PKs are dominated by a fast and slow clearance rate.

### S2 Initial Conditions

Initial conditions for the microglia equations were taken as the mean baseline microglia densities from PBS-treated control mice reported in [1], reflecting homeostatic densities in the OPL, IPL, and GCL/NFL. These initial conditions were used throughout all simulations except the rhesus macaque scenario shown in Figure 8 of the main text.

For the rhesus macaque data, initial conditions were likewise taken as the mean baseline microglia densities. However, only measurements from the peripheral retina were used, rather than those from the fovea or macula. The peripheral region represents $\geq 90\%$ of the total retinal surface area and is therefore most representative of the region likely to be captured in a randomly selected field-of-view image, consistent with the sampling approach used in [1]. Both the mouse and rhesus macaque initial conditions are provided in Table S1.

The initial peritoneal LPS concentration was determined from reported LPS doses, which were provided in units of mg/kg. To express this dose in ng/mL for the peritoneal compartment, the average body weight of the relevant experimental animals was inferred, and the corresponding mass of LPS delivered per mouse was divided by an approximate peritoneal fluid volume to obtain an initial concentration.

For the 9 mg per kg of body weight dose used in Dando *et al.*, the mean BALB/c mouse body weight of 24.7g (SDev = 1.83g) was used, which corresponds to an injected LPS mass of 0.2223 mg. To reflect intraperitoneal (IP) injection, the injected mass was divided by an approximate peritoneal fluid volume of 0.11 mL, giving

$$R(0) = \frac{0.2223 \text{ mg}}{0.11 \text{ mL}} = 2.021 \text{ mg/mL} = 2{,}021{,}000 \text{ ng/mL}.$$

For intravenous (IV) injection in mice, the same injected LPS mass was instead distributed within the systemic blood volume. Using a blood volume of 1.482 mL for a mouse of this body weight, the corresponding initial blood concentration was

$$B(0) = \frac{0.2223\,\text{mg}}{1.482\,\text{mL}} = 0.150\,\text{mg/mL} = 150{,}000\,\text{ng/mL}.$$

Analogously, for rat simulations, the initial peritoneal concentration was calculated as follows. Peritoneal fluid volume was taken as 3.07 mL [2] and rat weight was taken as 283 g [3], reflecting the mean body weight of the rats this subsystem was calibrated to. This corresponds to an injected LPS mass of 2.547 mg, yielding the initial peritoneal concentration

$$R(0) = \frac{2.547 \text{ mg}}{3.07 \text{ mL}} = 0.83 \text{ mg/mL} = 830,000 \text{ ng/mL}.$$

For rhesus monkey simulations, LPS was administered intravenously. A representative body weight of 6.75 kg, corresponding to the midpoint of the reported experimental range [4], was used. At a dose of 9 mg/kg, this yields a total administered LPS mass of 60.75 mg. Blood volume was estimated as 54 mL/kg of body weight [5], giving a total systemic blood volume of 364.5 mL. The corresponding initial blood concentration was therefore

$$B(0) = \frac{60.75 \text{ mg}}{364.5 \text{ mL}} = 0.1667 \text{ mg/mL} = 166,700 \text{ ng/mL}.$$

Circulating LPS was assumed to be absent prior to injection in all cases. All initial conditions used across experimental scenarios are summarized in Table S1.

#### S3 Alternative Models Tested Against the Microglia Dataset

The model presented in **Equations 4-6** in the main text was the result of many iterations. We considered several simpler versions of the model and fit them to the microglia density in the retina, i.e., the data in **Figure 3** of the main text. For the most part, we were unable to capture the nonlinear nature of the microglia migration patterns with any of these alternate models. We have summarised all the models we tried below along with their AIC and justification for why they were not chosen as the final model in **Table S2**. We use the following formula to calculate the AIC:

$$AIC = n \ln\left(\frac{RSS}{n}\right) + 2k, \tag{S5}$$

**Table S1:** Initial conditions used for each experimental scenario. $R(0)$ and $B(0)$ denote the initial LPS concentrations in the peritoneal and blood compartments, respectively, with $R(0)$ used only for intraperitoneal administration and $B(0)$ used for both intraperitoneal and intravenous administration (where $R(t)$ is omitted). $O(0)$, $I(0)$, and $G(0)$ denote baseline microglia densities in the OPL, IPL, and GCL/NFL. IP and IV denote intraperitoneal and intravenous administration, respectively.

| **Experimental scenario** | ***R(0)*** **[ng/mL]** (Peritoneal LPS) | ***B(0)*** **[ng/mL]** (Blood LPS) | ***O(0)*** **[cells/mm$^2$]** (OPL Microglia) | ***I(0)*** **[cells/mm$^2$]** (IPL Microglia) | ***G(0)*** **[cells/mm$^2$]** (GCL/NFL Microglia) |
|---|---|---|---|---|---|
| IP mice | 2,021,000 | 0 | 159.575 | 165.8 | 93.525 |
| IV mice | — | 150,000 | 159.575 | 165.8 | 93.525 |
| IP rats | 830,000 | 0 | 159.575 | 165.8 | 93.525 |
| IV rhesus monkeys | — | 166,700 | 143.37 | 185.27 | 86.00 |

where $n$ is the number of data points, $k$ is the number of free parameters, and $RSS$ is the residual sum of squares. All fits were conducted with a multistart algorithm using least squares nonlinear fitting. The purpose of this summary is not to claim that a formal model selection procedure was conducted. This would require more rigorous testing, which we feel is beyond the scope of this work. For information on how to conduct model selection, please see [6–8].

**Table S2: Summary of alternate models.** Below we summarise the mathematical models that were considered as possible alternates to **Equations 4-6** in the main text. Variables are equivalent to those in the main text, with the addition of $M$ representing total microglia across all retinal layers. The AIC was calculated using **Equation S5**

| Model | RSS | AIC | Justification for model rejection |
|---|---|---|---|
| $\frac{dO}{dt} = rO\left(1 - \frac{O}{K}\right) + s_O B$<br>$\frac{dI}{dt} = rI\left(1 - \frac{I}{K}\right) + s_I B$<br>$\frac{dG}{dt} = rG\left(1 - \frac{G}{K}\right) + s_G B$ | 18,917 | 108.86 | All retinal layers exhibited both increases and decreases in microglia counts during 96 hours, which contradicted the mathematical model. Furthermore, the RSS was large. |
| $\frac{dO}{dt} = -k_O O + k_{IO} I$<br>$\frac{dI}{dt} = k_O O - (k_{IO} + k_{IG})I + k_G G$<br>$\frac{dG}{dt} = -k_G G + k_{IO} I$ | 3,187 | 85.49 | The model was unable to fit the trend of the data, for example it was unable to capture the increase and then decrease in the OPL, and the RSS was still quite high. |
| $\frac{dO}{dt} = r_O OB - k_R O + k_{IO} I$<br>$\frac{dI}{dt} = r_I IB + k_O O - (k_{IO} + k_{IG})I + k_G G$<br>$\frac{dG}{dt} = r_G GB - k_G G + k_{IO} I$ | 98,805 | 118.19 | This model performed poorly and was unable to capture the small range of changes in each retinal compartment. |
| $\frac{dO}{dt} = \frac{rOB}{B + \eta} - \delta O$ | 183.87 | 21.32 | Fitting only the OPL compartment to this model with the Michaelis-Menten term provided an excellent fit to all measurements except 6 hours. |

| Model | RSS | AIC | Justification for model rejection |
|---|---|---|---|
| $\frac{\mathrm{d}O}{\mathrm{d}t} = -(k_{OG} + k_{OI})O + k_{IO}I + k_{GO}G$<br>$\frac{\mathrm{d}I}{\mathrm{d}t} = -(k_{IG} + k_{IO})I + k_{OI}O + k_{GI}G$<br>$\frac{\mathrm{d}G}{\mathrm{d}t} = -(k_{GO} + k_{GI})G + k_{OG}O + k_{IG}I$ | 2,090 | 73.92 | While the model fits the data better, this model doesn't depend on the LPS concentration in the blood. It had similar issues with its trend to the second model above. |
| $\frac{\mathrm{d}O}{\mathrm{d}t} = -k_{OI}O + k_{IO}I + k_{BO}B$<br>$\frac{\mathrm{d}I}{\mathrm{d}t} = -(k_{IG} + k_{IO})I + k_{OI}O + k_{GI}G$<br>$\frac{\mathrm{d}G}{\mathrm{d}t} = -(k_{GO} + k_{GI})G + k_{OG}O + k_{IG}I$ | 1,788 | 74.58 | The fitted curve matched the general trend of the data in each retinal layer but was unable to fit the data points and most importantly missed the data point at 6 hours in the OPL compartment. |
| $\frac{\mathrm{d}O}{\mathrm{d}t} = -k_{OI}OB_G + k_{IO}B_OI$<br>$\frac{\mathrm{d}I}{\mathrm{d}t} = k_{OI}OB_G - k_{IO}B_OI - k_{IG}IB_G + k_GG$<br>$\frac{\mathrm{d}G}{\mathrm{d}t} = k_{IG}IB_G - k_GG$<br>$\frac{\mathrm{d}R}{\mathrm{d}t} = -k_RR$<br>$\frac{\mathrm{d}B_O}{\mathrm{d}t} = f_OFk_R\left(\frac{Vol_P}{Vol_B}\right)R - k_BB_O$<br>$\frac{\mathrm{d}B_G}{\mathrm{d}t} = (1 - f_O)Fk_RB\left(\frac{Vol_P}{Vol_B}\right)R - k_BB_G$ | 1,859 | 70.51 | In this model, we tested the hypothesis that there were two different effects of LPS driving the microglia migration: one on the OPL compartment and one on the GCL compartment. This had a slight improvement over the previous models but introduced additional parameters. |
| $\frac{\mathrm{d}O}{\mathrm{d}t} = -\frac{k_{OI}OB_G}{B_G + \eta} + \frac{k_{IO}IB_O}{B_O + \eta}$<br>$\frac{\mathrm{d}I}{\mathrm{d}t} = \frac{k_{OI}OB_G}{B_G + \eta} - \frac{k_{IO}IB_O}{B_O + \eta} - \frac{k_{IG}IB_G}{B_G + \eta} + k_GG$<br>$\frac{\mathrm{d}G}{\mathrm{d}t} = \frac{k_{IG}IB_G}{B_G + \eta} - k_GG$<br>$\frac{\mathrm{d}R}{\mathrm{d}t} = -k_RR$<br>$\frac{\mathrm{d}B_O}{\mathrm{d}t} = f_OFk_R\left(\frac{Vol_P}{Vol_B}\right)R - k_BB_O$<br>$\frac{\mathrm{d}B_G}{\mathrm{d}t} = (1 - f_O)Fk_RB\left(\frac{Vol_P}{Vol_B}\right)R - k_BB_G$ | 577.855 | 58.49 | This model fit the data really well and captured all trends and data points. However, as we didn't have measurements for $B_O$ and $B_G$, the model was not practically identifiable for $f_O$. We also felt there was less biological justification for splitting $B$ into $B_O$ and $B_G$. |
| $\frac{\mathrm{d}O}{\mathrm{d}t} = -\frac{k_{OI}OB}{B + \eta} + \frac{k_{IO}IB}{B + \eta}$<br>$\frac{\mathrm{d}I}{\mathrm{d}t} = \frac{k_{OI}OB}{B + \eta} - \frac{k_{IO}IB}{B + \eta} - \frac{k_{IG}IB}{B + \eta} + k_GG$<br>$\frac{\mathrm{d}G}{\mathrm{d}t} = \frac{k_{IG}IB}{B + \eta} - k_GG$ | 1,220 | 67.46 | This model fit the IPL and GCL compartments well, but by removing the breakdown of $B$ into $B_G$ and $B_O$ we no longer fit the OPL compartment. |

#### S4 Estimating the Half-effect Constant $\eta_O$

As mentioned in the main text, $\eta_O$ and $\eta_G$ were not both practically identifiable with the data we had available. As such, we fixed $\eta_O$ with a value informed by our biological understanding of the experimental system. Experimental studies span a broad range of administered doses for LPS (approximately 0.1–9 mg/kg), from low-dose systemic challenges (0.1 mg/kg) to commonly used neuro-inflammatory paradigms at 4–5 mg/kg and higher-dose retinal inflammatory models (9 mg/kg) [1, 9–11]. For example, Michels *et al.* [11] used a final concentration of 5 mg/kg in their study of the effect of LPS on microglia. This value was determined after they conducted a dose-finding study prior to their experiment. Qin *et al.* [9] also injected mice intraperitoneally with a single dose of LPS of 5 mg/kg, the dosage of which was based on their previous study of endotoxic shock [12].

Other studies that did not focus necessarily on neuroinflammation used other dosages. For example, Alheim *et al.* [13] used 0.1 mg/kg in their experiments but were focused on the murine peripheral immune response, as opposed to generating a response in the central nervous system. Kozak *et al.* [14, 15] investigated the safety of a range of dosages for eliciting a fever in mice that the mice were able to recover from in a short period of time. They chose 1, 2.5 and 3 mg/kg. Studies examining large dosages have shown that fatality in mice is seen for 20 mg/kg [16]; however, this does not rule out fatality for concentrations below this.

Previous research by Banks *et al.* commented on the minimal penetration across the BBB murine barrier of LPS [10]. As a result of this, to estimate $\eta_O$, we chose to simulate an injection of LPS at 4 mg/kg as both a pragmatic intermediate value and a literature standard that induces a clear but non-saturating neuroinflammatory response. Simulating 4 mg/kg we estimated $\eta_O$ as the peak concentration in the blood experienced from this injection, i.e. $\eta_O = 1667$ ng/mL, see Figure S1. As we are estimating the half-effect in the OPL compartment, we expect this to be much higher than the GCL/NFL compartment as the OPL layer is considerably further from the vasculature.

To confirm this as a realistic choice for $\eta_O$, we investigated values in the literature for *in vitro* studies of LPS and their effect on microglia and other innate immune cells. Kumar *et al.* [17] and Chang *et al.* [18] noted that a concentration of 500 ng/mL of LPS induced the maximum level of nitrite from BV2 microglia with no significant cytotoxicity. Whereas Rossi-George *et al.* [19] commented that maximal release from BV2 cells was seen from 1000 ng/mL. The concentration of 1000 ng/mL has also been used by Pan *et al.* to elicit a response from BV2 cells [20].

We also obtained EC50 and IC50 LPS measurements for IL-6 released from macrophages [21] and microglia cell viability [22], see Figure S2, using the following equations:

$$E = \frac{(E_{max} - E_{min})C^h}{C^h + EC_{50}^h} + E_{min}, \qquad I = \frac{(I_{max} - I_{min})IC_{50}^h}{C^h + IC_{50}^h} + I_{min},$$

$E$ and $I$ are the effect functions for the dose-response curves and $C$ is the concentration of LPS. The maximum and minimum effects are $E_{max}, I_{max}$ and $E_{min}, I_{min}$ respectively, and $h$ is the hill coefficient. Interestingly, the IC50 we obtained for cell viability of microglia exposed to different LPS dosages was very close to our chosen value of $\eta_O$, whereas the EC50 obtained for IL-6 secretion from macrophages was very close to our fitted value of $\eta_G$ (see the main text).

Overall, we are confident that $\eta_O$ represents a biologically resonable choice. It is possible this value could vary and with more data it may be possible to estimate it with more certainty.

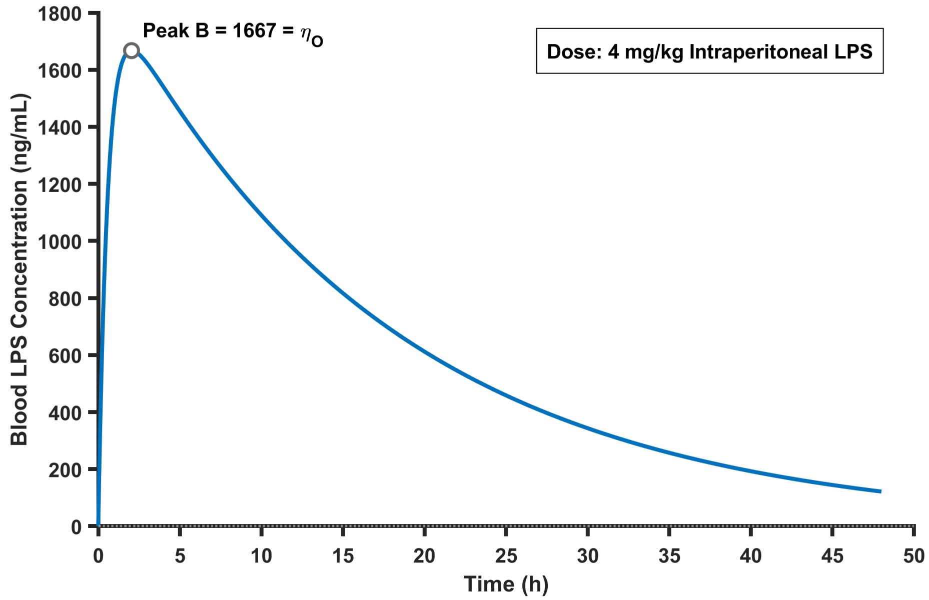


**Figure S1**: **Estimation of the half-activation constant $\eta_O$ from an intermediate LPS dose.** Blood LPS concentration $B(t)$ simulated using the intraperitoneal pharmacokinetic subsystem at a representative intermediate dose of 4 mg/kg. The peak concentration ($\approx$ 1667 ng/mL) is highlighted and taken as a proxy for the half-activation constant $\eta_O$, corresponding to the LPS level required to induce a half-maximal microglia migratory response toward the OPL. This approach assumes that an intermediate, non-saturating inflammatory stimulus provides a biologically reasonable estimate of the activation threshold in the absence of direct dose–response measurements.

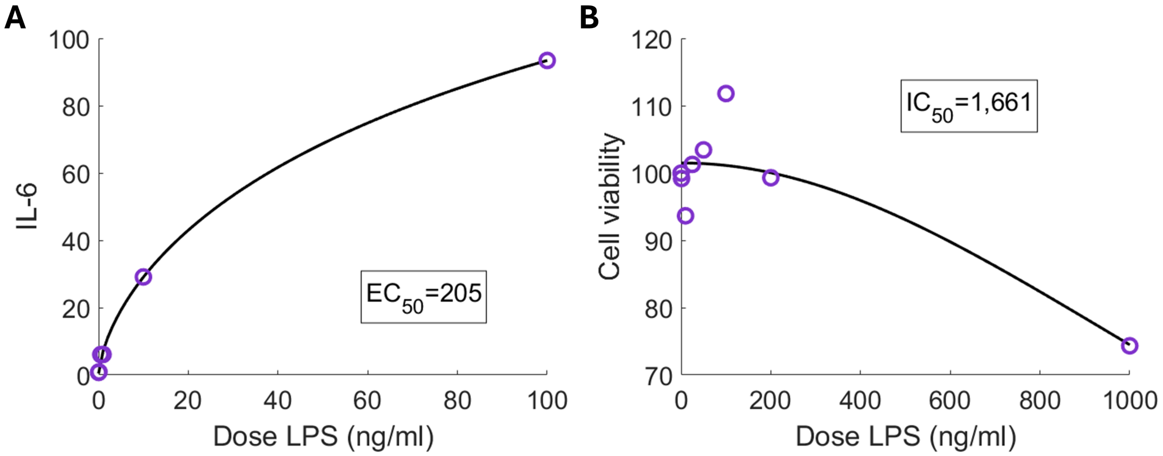


**Figure S2**: **Fitting dose-response curves to data to obtain an EC50.** (**A**) IL-6 production from macrophages was measured for a range of LPS dosages [21]. Fitting a dose-response curve to this data produced an EC50 of 205 ng/mL. (**B**) Cell viability of microglia were measured for a range of LPS dosages [22]. Fitting a dose-reponse curve to this data gave an IC50 of 1,661 ng/mL.

### S5 Supplementary Results

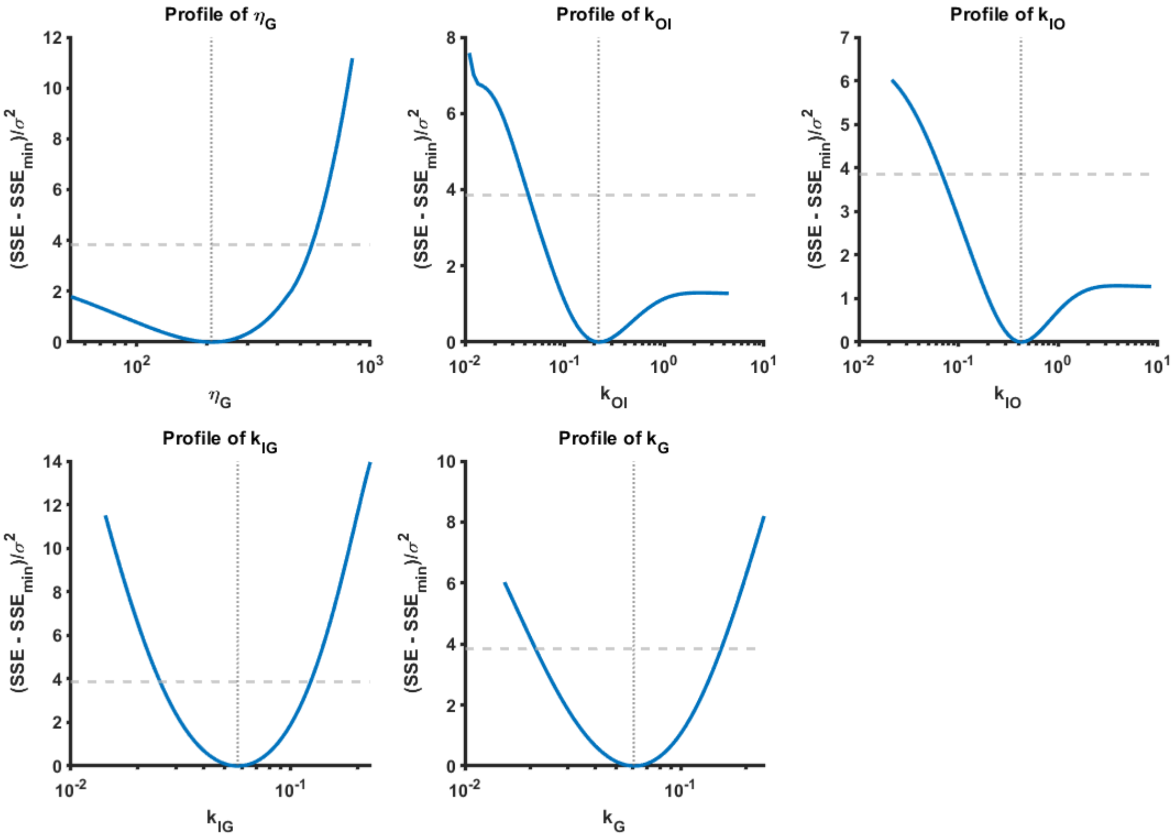


**Figure S3**: **Profile likelihoods for fitted microglia migration parameters.** Profile likelihood curves for the parameters estimated by nonlinear least-squares fitting of the microglia migration subsystem to the retinal microglia density data of Dando *et al.* [1]. Each panel shows the change in model error as a single parameter is varied while all others are re-fitted. The dashed horizontal line represents the 95% likelihood-ratio confidence threshold ($\chi^2_{0.95,1}$ = 3.84) used for assessing practical identifiability. Distinct minima are observed for each parameter, meaning that the microglia migration parameters are practically identifiable from the available data.

**Table S3: Fitting statistics for the microglia migration model following IP LPS administration.** RMSE values correspond to the model fits shown in Figure 4D–F of the main text and quantify the typical discrepancy between model predictions and experimental measurements. RMSE is reported in units of cells/mm$^2$.

| **Retinal compartment** | **RMSE** |
|---|---|
| Outer plexiform layer (OPL) | 6.54 |
| Inner plexiform layer (IPL) | 8.48 |
| Ganglion cell & nerve fibre layer (GCL & NFL) | 7.73 |

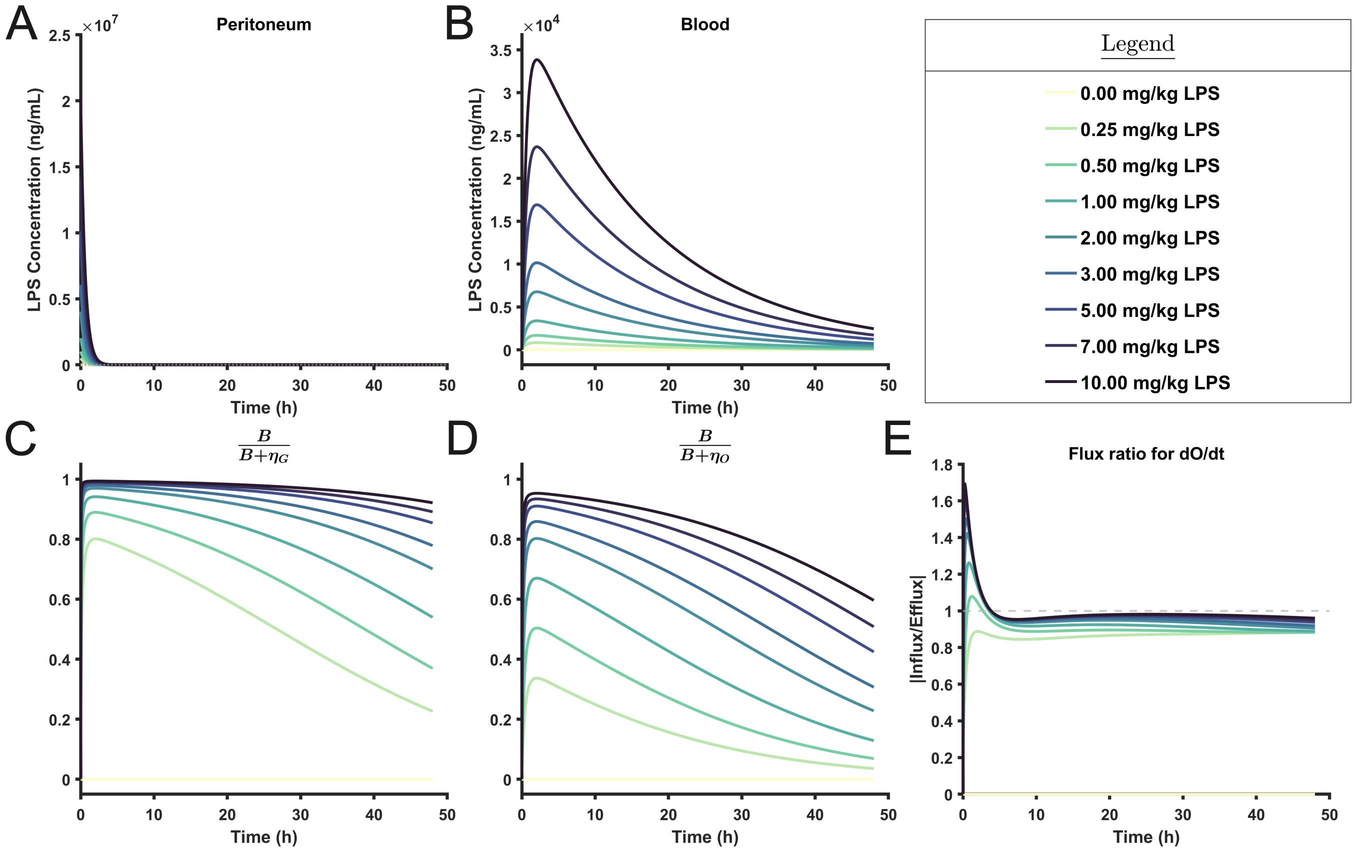


**Figure S4**: **Additional model variables during intraperitoneal LPS dose escalation.** Outputs from the dosage sweep of the coupled PK–microglia migration model. Each panel shows auxiliary variables used to aid interpretation of the dose–response behaviors presented in the main text. (**A**) Peritoneal LPS concentration trajectories corresponding to each simulated dose. (**B**) Resulting blood LPS concentrations. (**C**) Hill activation term regulating migration toward the GCL/NFL. (**D**) Hill activation term regulating migration toward the OPL. (**E**) Ratio of influx to efflux in the OPL equation, included to clarify the nonmonotonic OPL dose–response behavior observed in Figure 4A. All simulations were performed using the fixed parameter regime provided in Table 3 of the main text, varying only the initial dosage.

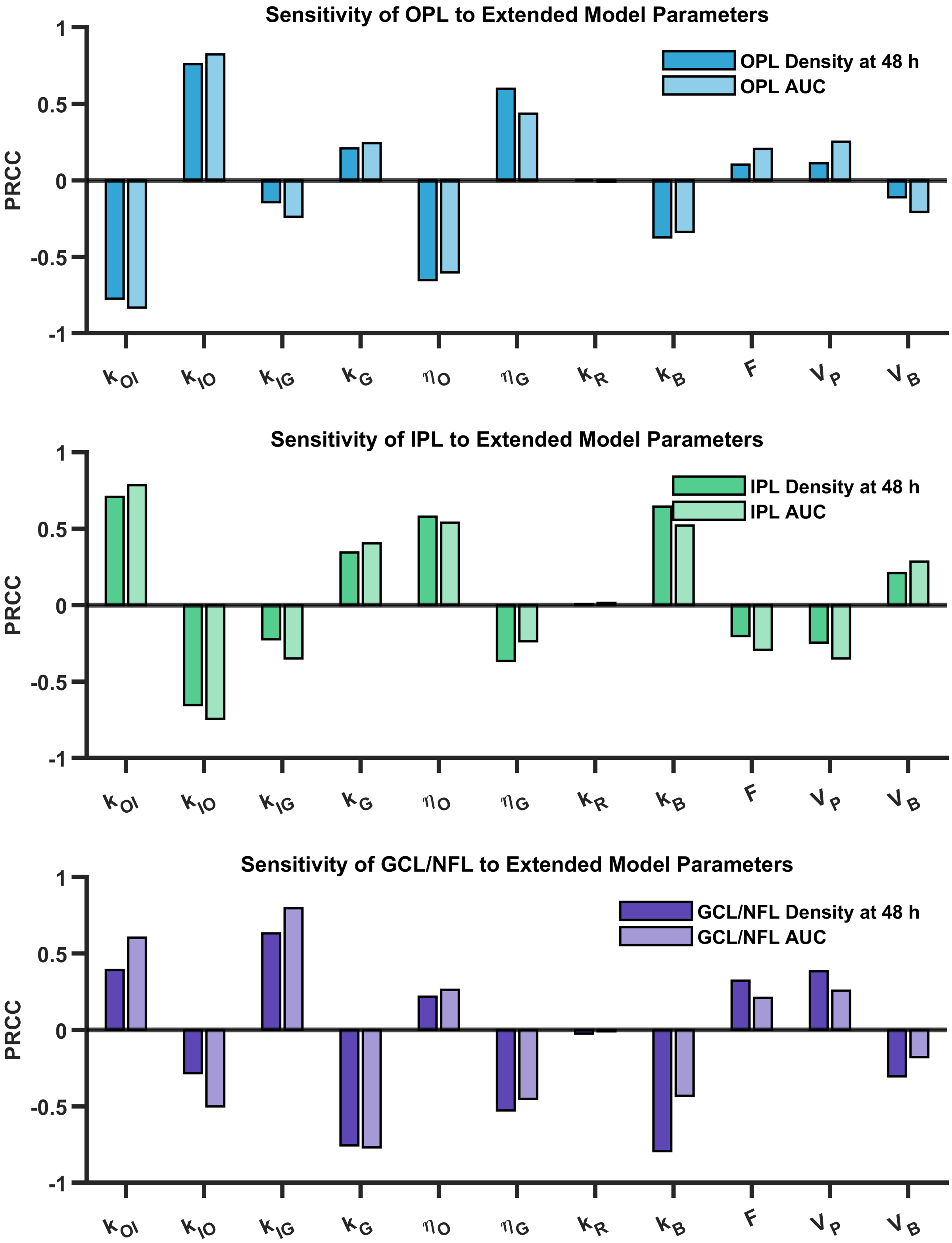


**Figure S5**: **Extended global sensitivity analysis using widened parameter ranges.** Partial rank correlation coefficients (PRCCs) were computed in the same manner as in Figure 5 of the main text, but using expanded parameter ranges. Each parameter was allowed to vary from one third of its original lower bound up to three times its original upper bound in order to explore model behavior under extreme conditions. (top row) Sensitivity of OPL microglia density outcomes for final density at 48h and the 0–48h area under the curve (AUC). (middle row) As in (top row), but for IPL microglia density and IPL AUC. (bottom row)As in (top row), but for GCL/NFL microglia density and GCL/NFL AUC.

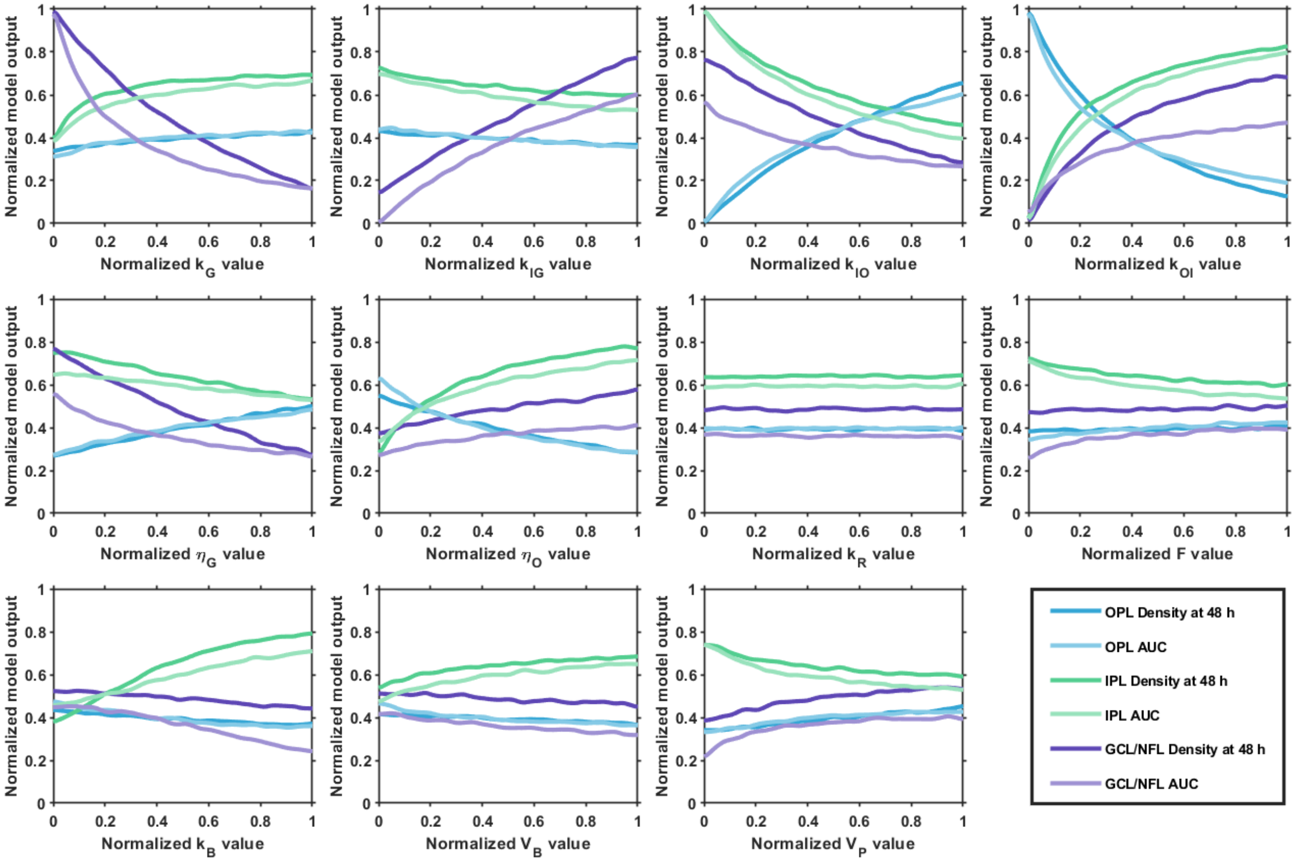


**Figure S6**: **Monotonicity assessment for partial rank correlation coefficient analysis.** Normalized model outputs are plotted against each parameter to assess whether the relationship between that parameter and the six microglia migration outcomes is predominantly monotonic. Each panel shows the response of the six outcomes (48h density and 0–48h AUC for the OPL, IPL, and GCL/NFL) across the full range of each parameter while the others vary simultaneously. Overall trends are monotonic for all parameters, satisfying the monotonicity requirement for the use of partial rank correlation coefficients. Small deviations from strict monotonicity appear, though this is expected as a consequence of varying all parameters simultaneously and does not affect the interpretability of PRCCs.

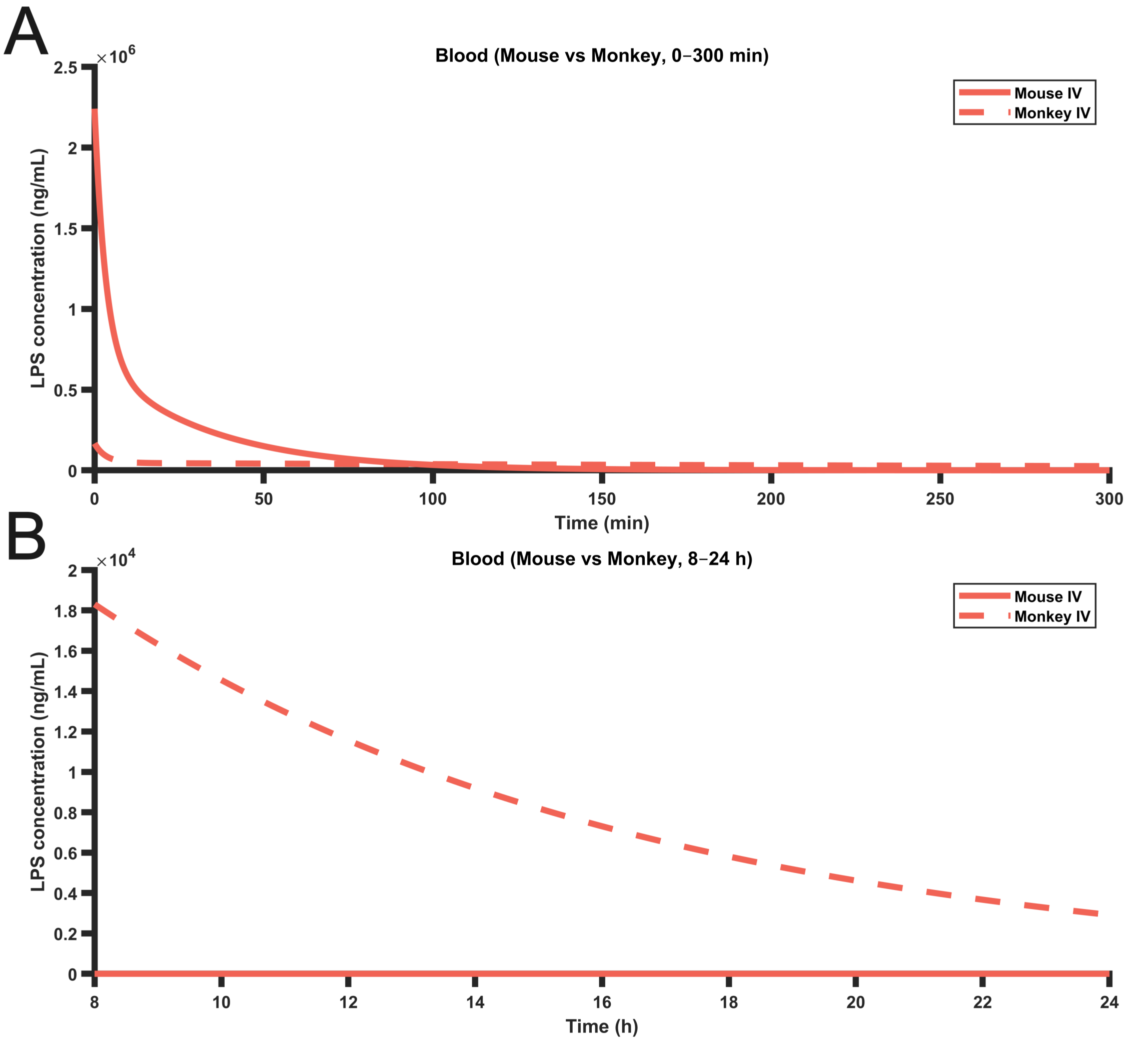


**Figure S7**: **Prolonged microglia migration in rhesus monkeys arises from sustained blood LPS despite lower early-time concentrations.** Blood LPS concentration profiles for the mouse and monkey models over two time windows. (A) The first 300 minutes post-treatment, during which the monkey model exhibits substantially lower LPS availability than the mouse model. (B) the later 8–24 hours post-treatment, where the rhesus monkey model demonstrates comparatively higher LPS availability. Together, these panels explain why the monkey model exhibits a longer-lasting microglia response in Figure 8C of the main text.

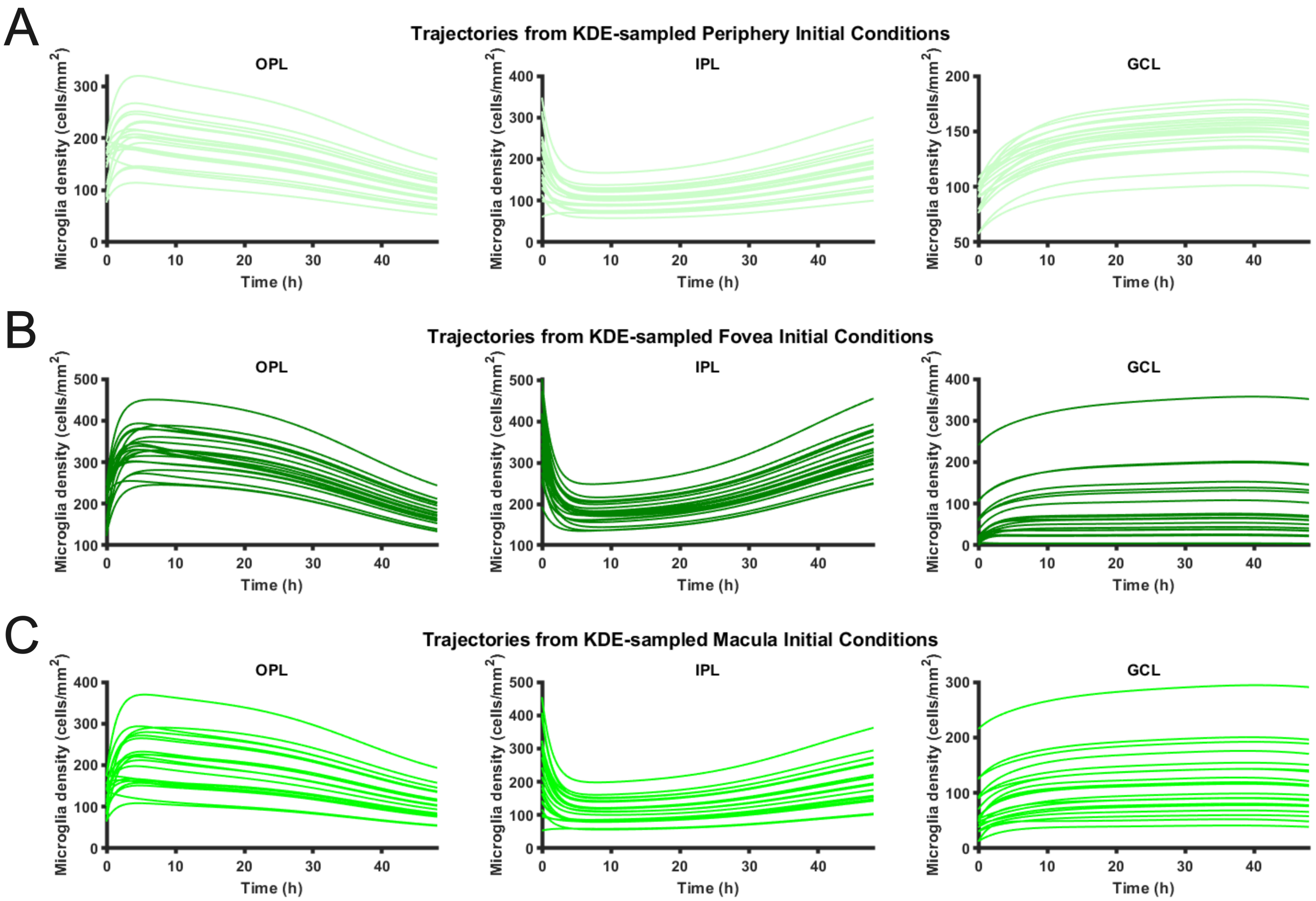


**Figure S8**: **Propagation through the model smooths heterogeneous initial microglia distributions into unimodal outcomes.** Twenty model trajectories generated by simulating the rhesus monkey model across experimentally derived initial microglia distributions. Panels (**A**–**C**) show ensembles of microglia density trajectories in the OPL, IPL, and GCL/NFL arising from kernel-density–sampled initial conditions from the retinal periphery, fovea, and macula, respectively. Despite heterogeneity and bimodality in the initial distributions (Figure 9B), trajectories rapidly converge toward a narrower family of responses within each retinal layer.
